# HMCES corrupts replication fork stability during base excision repair in homologous recombination deficient cells

**DOI:** 10.1101/2024.07.31.605977

**Authors:** María José Peña-Gómez, Yaiza Rodríguez-Martin, Marta del Rio Oliva, Jean Yves Masson, José Carlos Reyes, Iván V. Rosado

## Abstract

Apurinic/apyrimidinic (AP) sites and single-strand breaks (SSB) arising from base excision repair (BER) during misincorporation of damaged nucleobases may hinder replication fork stability in homologous recombination-deficient (HRD) cells. At templated AP-sites, HMCES DNA-protein crosslinks (DPC) regulate replication fork speed while avoiding APE1-mediated cytotoxic double-strand breaks (DSB). Whereas the role of HMCES at template DNA strand is well studied, its consequences on nascent DNA are less understood. Here, we provide evidence that HMCES play detrimental roles during removal of 5- hydroxymethyl-2’-deoxycytidine (5hmdC)-derived 5-hydroxymethyl-2’-deoxyuridine (5hmdU) by BER at replication forks. HRD cells display heightened HMCES chromatin levels, which increase upon 5hmdC exposure, suggesting that HMCES binds both spontaneous and 5hmdC-induced AP-sites. HMCES depletion largely suppresses 5hmdC- mediated *Fancd2*^-/-^ replication fork defects, chromosomal aberrations and cell lethality, suggesting that HMCES is responsible for the replication fork impairment and lethality observed in HRD cells. Therefore, HMCES-DPCs are a novel source of BER-initiated PRIMPOL-mediated ssDNA gaps, implying endogenous DPCs as a source of DNA damage in HRD tumours.

**Teaser:** Covalent binding of HMCES to nascent DNA blocks replication progression and kills homologous recombination deficient cancer cells

## Introduction

Fanconi anaemia (FA) is an ultrarare genetic syndrome mainly characterized by congenital abnormalities, progressive bone marrow failure due to aldehyde-dependent depletion of hematopoietic stem cells (HSC) (*1, 2*), and extreme cancer susceptibility (*3, 4*). Upon DNA damage, stalling of DNA replication leads to ATR-checkpoint kinase activation to facilitate FA-core complex dependent monoubiquitylation of the clamped FANCI/FANCD2 (ID2) heterodimer recruited around sites of DNA lesions(*5, 6*). Ubiquitylated ID2 complex recruits and stabilizes critical components of the DNA damage response (DDR), such as homologous recombination (HR) repair factors like FAN1, SLX4, XPF, BRCA1/2 and RAD51 to promote replication fork protection or the classical interstrand crosslink (ICL) repair mechanism(*7, 8*). Heightened chromosomal aberrations caused by defects in the FA pathway lead to extreme hypersensitivity to endogenous reactive aldehydes(*2, 9, 10*) or exogenous ICL-inducing compounds such as cisplatin, mitomycin C (MMC) or diepoxybutane (DEB)(*11*), which are typically used in the clinic as diagnostic tool(*12*). In addition to their role during ICL-repair, some FA gene products play crucial functions in other DNA transactions processes such as DNA double-strand breaks (DSBs)(*13*) or DNA replication forks protection during perturbation of DNA synthesis thus avoiding replication stress (RS) (*14–16*). Frequent sources of RS include exposure to exogenous agents used therapeutically such as the dNTP-depleting compound hydroxyurea (HU), MMC, the PARP-trapping inhibitor Olaparib or the endogenous formation of DNA-protein crosslinks (DPCs), thus contributing to genomic instability in cancer cells(*17–21*). Therefore, a deeper understanding of the endogenous and exogenous sources that trigger RS and the molecular mechanisms contributing to accurate maintenance of the genomic integrity are of paramount relevance.

Base depurination and misincorporation of damaged nucleobases such as oxidized guanines, thymine analogues and deamination derivatives such as 5’-hydroxymethyl-2’-deoxyuridine (5hmdU), 5’-formyl-2’-deoxyuridine (5fdU), 5’-carboxy-2’-deoxyuridine (5cadU) are amongst the commonest spontaneous base lesions in DNA(*22–25*). These adducted bases efficiently trigger canonical base excision repair (BER) (*26, 27*). During the removal of the damaged base, DNA glycosylases generate apurinic/apyrimidinic (AP) sites, which are enzymatically processed to DNA single-strand breaks (SSBs) by AP endonuclease 1 (APE1). Upon DNA end processing by PNKP or APTX, POLβ or POLλ conducts gap filling and the XRCC1/LIG1 complex or LIG3 seals the nicked DNA(*28–30*). Despite the “pass the baton” BER model, under certain conditions BER intermediates such as AP sites or SSBs may persist and challenge DNA replication causing RS. During the RS response, several protective mechanisms are employed to minimize ssDNA gaps accrual(*19, 31–33*). Uncoupling of the CMG helicase and DNA polymerases results in persistence of single-stranded DNA (ssDNA) stretches rapidly activating the RPA-ATR-CHK1 axis, which leads to cell cycle arrest while DNA repair occurs. A mechanism to avoid genomic instability is the process of replication fork reversal (RFR) (*34*), which gives cells time enough to resolve damaged DNA ahead the fork and subsequent resumption of DNA synthesis. An alternative mechanism termed “repriming” involving the recently described activities of the primase-polymerase (PRIMPOL) has also been reported(*35–37*). Upon fork stalling, PRIMPOL reprimes at the leading strand ahead of both the stalled replication fork and the lesion, to complete DNA synthesis leaving of translesion synthesis (TLS) polymerases REV1-POLζ-POLQ, belonging to the DNA damage tolerance (DDT) pathways(*38–40*). Alternatively, the error-free template-switching (TS) recombination mechanism may also occur under certain conditions. These replication fork rearrangements take place to complete DNA synthesis at the expenses of ssDNA gaps formation whilst minimizing genetic instability throughout postreplicative repair. Therefore, delayed or persistent SSBs at nascent DNA strands have been described to account for exquisitely cytotoxicity to poly-(ADP-ribose) polymerase (PARP) trapping drugs such as Olaparib observed in HRD cells (*17, 41*). However, the underlying molecular basis of this cytotoxicity still remains elusive. Several mechanisms have been recently proposed as major contributors of HRD cell lethality, of which unifying feature is the binding and trapping of PARP1 to unprocessed ssDNA gaps behind the fork (*20, 42*). Recently, 5hmdU or 2’-deoxyuridine (dU) misincorporation has emerged as a novel source of replication fork instability (*15, 43–45*). Replication fork protection and gap suppression separation-of-functions BRCA2 mutant cells have been reported to sensitize at similar extent to PARP inhibition(*43*). As replication forks misincorporate 5hmdU, BER activity is predicted to generate repair intermediates resulting in replication fork-BER intermediate collisions accounting for HRD cell lethality (*15, 43–45*). Whereas mechanisms accounting for ssDNA gaps formation at or behind the replication forks induced by HU or other chemotherapeutic agents are well documented (*20, 35, 42, 46, 47*), the mechanism by which BER-mediated nascent ssDNA gaps arises is currently under active investigation. If dU or 5CldU misincorporation remains unrepaired throughout the cell cycle, these aberrant bases in template DNA activate the ATR signaling and induce replication fork collapse (*48, 49*). However, their molecular consequences during the current cell cycle are barely unexplored(*15, 44*).

HMCES has been previously demonstrated to associate to replication forks and covalently binds to AP sites throughout a thiazolidine bond between the Cys2 residue and the aldehydic form of AP sites, forming a HMCES-DPC on ssDNA(*50–53*). HMCES has higher binding affinity for ssDNA over double-stranded DNA (dsDNA) (*50, 52*). Upon formation of HMCES-DPC on template DNA strand, replication fork speed decreases and APE1-mediated cytotoxic DSB at the fork is avoided(*50*). Lesion bypass is assisted by the FANCJ denaturation activity of HMCES(*54*), and subsequent TLS synthesis, thus protecting cells from AP site genotoxicity. Then, HMCES is removed from DNA by at least two proposed mechanisms, self-reversal catalyzed by the Glu127 residue(*55, 56*), or degradation in a SPTRN and proteasome-mediated fashion(*50, 54*). Whereas AP site protection at template DNA by HMCES is well studied, the molecular and cellular consequences of HMCES binding to nascent DNA is less understood.

Here, we provide evidence that HMCES binding to AP sites in nascent DNA strand plays a detrimental role during removal of 5hmdU by BER at replication forks. HRD cells display heightened chromatin levels of HMCES, which increase upon 5-hydroxymethyl-2’-deoxycytidine (5hmdC) exposure, suggesting that HMCES binds to spontaneous and 5hmdU-induced AP sites. HMCES is responsible for replication fork impairment, genomic instability and cytotoxic phenotypes observed in HRD cells during replication fork-associated 5hmdU BER. Genetic depletion of SMUG1 or APE1/2 BER factors, which generate ssDNA gaps, largely suppress 5hmdU-mediated phenotypes observed in HRD cells. However, loss of downstream ssDNA gap filling factors PARP1 or XRCC1, impacts on the persistence of ssDNA gaps and exacerbated these phenotypes. Moreover, depletion of HMCES largely suppresses 5hmdC-mediated replication fork defects, chromosomal aberrations and cell lethality in HRD cells. These data suggest that HMCES-DPCs formed on nascent DNA AP sites upon removal of misincorporated 5hmdU are responsible for the replication fork defects observed in HRD-deficient cells, and place endogenous DPCs as a novel source of genomic instability in HRD tumors.

## Results

### Misincorporation of 5hmdU arising from deaminated 5hmdC causes genomic instability in HRD cells

Previous data from our group and others have shown that HRD cells lacking FANCD2, MUS81, BRCA1 or BRCA2 were sensitive to cytotoxic misincorporation of 5hmdU on genomic DNA. 5hmdU can arise from either the 5hmdC salvage pathway or the cytosine demethylation process(*15, 43, 45*). To address the molecular mechanisms accounting for the cellular genotoxicity of 5hmdU- mediated BER under inactivation of the replication fork maintenance FA/BRCA pathway, we sought to investigate accumulation of ssDNA gaps as main driving source of genomic instability. We firstly examined whether loss of genes involved in the cytidine salvage pathway, deoxycytidine kinase *Dck* or dCMP deaminase *Dctd*, had any effect on the viability of 5hmdC-treated *Fancd2^-/-^* MEFs. Consistent with previously results in human HRD cells (*15, 45*), knockdown of either *Dck* or *Dctd* largely suppressed 5hmdC-mediated *Fancd2^-/-^* lethality whereas viability of *wild type* cells remained unaltered **(Figure 1A and supplementary figure 1)**, suggesting that 5hmdU arising from cytosolic 5hmdC deamination mediates HRD cell cytotoxicity and recapitulates previous data obtained in human cells. In the absence of the FA/BRCA protection pathway, BER intermediates of misincorporated 5hmdU on nascent DNA, triggered a DDR and impaired replication fork protection and chromosomal stability. Consistent with this, Olaparib-mediated PARP trapping or CHK1 inhibition by UCN-01 or AZD7762 exacerbated 5hmdC-mediated DDR signalling and chromosome instability in *Fancd2^-/-^* cell, suggesting that ssDNA gaps arising during BER-mediated 5hmdU removal were signalled by PARP1 and probably accounted for the chromosomal instability and cytotoxic phenotypes observed in *Fancd2^-/-^*cells(*15*). Moreover, ssDNA gaps accumulation during S-phase due to chemical or genetic perturbation of Okazaki fragments maturation (OFM) caused extensive PARP1-dependent nuclear PARylation(*57, 58*). We therefore sought to determine if 5hmdC treatment affected nuclear PARylation in total or EdU^+^ cell population, and SSBs in HRD cells. As expected, *Fancd2^-/-^* EdU^+^ cells displayed heightened nuclear PARylation levels upon 5hmdC treatment **(Figure 1B)**, also confirmed in total nuclear PARylation levels **(supplementary figure 2A)**, while remained relatively unchanged in *wild type* cells. Heightened PARylation was largely dependent on PARP1, as PARP1 knockdown significantly decreased 5hmdC-induced nuclear PARylation in *Fancd2*^-/-^ cells while remained unaffected in *wild type* cells **(Figure 1B and supplementary figure 2A)**. As an indirect measurement of SSBs associated to replication forks, we determined PAR levels associated to ongoing replication forks by “*in situ* protein interaction with nascent DNA replication forks” or SIRF assay. Compared to *wild type* MEFs, *Fancd2*^-/-^ cells displayed heightened EdU-PAR foci, which were further exacerbated by 5hmdC treatment **(Figure 1C)**. This increase in EdU-PAR SIRF foci associated to nascent DNA observed in *Fancd2*^-/-^ cells was largely dependent on PARP1, as PARP1 knockdown significantly reduced both spontaneous and 5hmdC-induced EdU-PAR foci **(Figure 1C)**, not reaching spontaneous levels, which suggest We next performed alkaline comet assay to assess the formation of SSBs or ssDNA gaps upon 5hmdC exposure in a direct manner. In agreement with heightened PARylation and EdU-PAR foci observed in *Fancd2^-/-^* cells, we observed a significant increase of spontaneous SSBs in *Fancd2^-/-^*MEFs **(Figure 1D)**. Moreover, whereas 5hmdC treatment almost had any effect on SSBs levels in *wild type* cells, a significant increase was detected in the absence of FANCD2 **(Figure 1D)**, suggesting that misincorporation of 5hmdU results in accumulation of ssDNA gaps in the absence of FANCD2, which probably accounts for the heightened PARylation levels at or in close proximity to replication forks. In agreement with this notion, we also found that 5hmdC exposure significantly affected the replication fork symmetry index in *Fancd2^-/-^* cells **(Figure 1E)** and increased the frequency of sister chromatids exchanges (SCEs) **(Figure 1F)**, however we did not observe any evidence of heightened ser345-CHK1 or ser33-RPA2 under these conditions **(supplementary figure 3)**, suggesting that 5hmdC exposure is not a strong signal to activate the DNA damage checkpoint in the absence of FANCD2. These results indicate that ssDNA gaps arising during removal of 5hmdU induces direct replication fork stalling in HRD cells and increases sister chromatid recombination.

**Figure 1.**
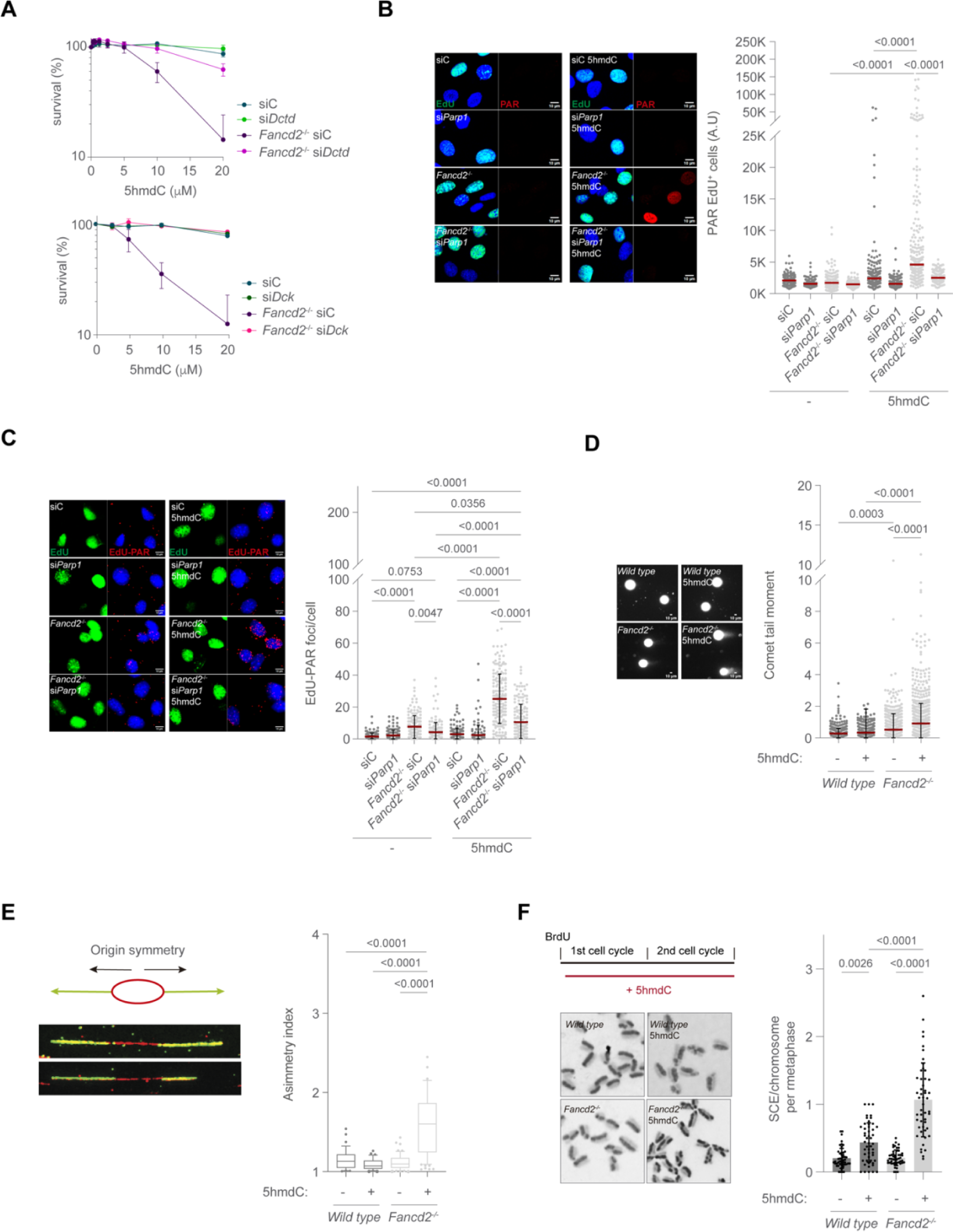
Genomic instability induced by 5hmdC in HRD cells A. MTT cell proliferation assay of *wild type* or *Fancd2*^-/-^ cells knocked down for *Dctd* (*upper plot*) or *Dck* (*lower plot*) exposed to the indicated dose of 5hmdC for 3 days. B. *Left*, representative PAR (red) immunofluorescence images of EdU^+^ (green) *wild type*, *Fancd2^-/-^*, si*Parp1* and *Fancd2^-/-^* si*Parp1* cells exposed to 5hmdC (10µM) for 3h. DAPI (blue) stains nuclear DNA. *Right*, plot depicting PAR mean intensity signal per nucleus. C. *Left*, representative images of EdU^+^-PAR foci (red) by SIRF analysis from *wild type*, *Fancd2^-/-^*, si*Parp1* and *Fancd2^-/-^*si*Parp1* cells exposed to 5hmdC (10µM) for 3h. DAPI (blue) stains nuclear DNA and EdU (green) stains S-phase cells. *Right*, plot depicting EdU-PAR foci per nucleus (n=2. D. *Left*, representative images of alkaline comet assay from *wild type* and *Fancd2^-/-^* cells following 5hmdC treatment (10µM) for 3h. *Right*, plot depicting comet tail moment per cell (n=4. E. *Top left*, scheme of the DNA fiber origin symmetry. *Bottom left*, representative images of DNA fiber from *wild type* and *Fancd2*^-/-^ cells after 5hmdC treatment (40µM) for 30 min. *Right*, box plot of the asymmetry index. F. *Top left*, scheme of SCE assay. *Bottom left*, representative images of SCEs from *wild type* and *Fancd2^-/-^* cells exposed to 5hmdC (10µM) for 40h. *Right*, bar plot of SCE/chromosome per metaphase (n=50 of each of 2 biological replicates; bar represents mean ± s.d.).

### ssDNA gaps arising from the concerted activities of SMUG1 and APE1/2 on misincorporated 5hmdU promote replication fork instability

Previous studies have reported that accumulation of ssDNA gaps is a major contributor of BRCA1/2 tumour cell death(*19, 20, 59*). Replication fork stalling elicits a primase-polymerase (PRIMPOL)- dependent repriming activity resulting in the formation of ssDNA gaps, which are subsequently filled in a POLθ or RAD18-REV1-POLζ dependent TLS manner (*38, 40*). As SMUG1 is the main DNA glycosylase removing 5hmdU from genomic DNA, we sought to determine the contribution of SMUG1 to the formation of ssDNA gaps in nascent DNA during BER-dependent 5hmdU removal. In agreement with previous data, SMUG1 knockdown had almost no effect in spontaneous nor 5hmdC-induced nuclear PARylation in *wild type* EdU^+^ cells. However, nuclear PARylation in SMUG1-depleted *Fancd2^-/-^* cells significantly decreased to untreated levels upon 5hmdC exposure in both total or EdU^+^ cells **(Figure 2A and supplementary figure 2B)**, suggesting that SMUG1 is responsible for most, if not all, 5hmdC-mediated nuclear PAR signal observed in *Fancd2*^-/-^ cells. Moreover, SMUG1 knockdown also suppressed 5hmdC-mediated EdU-PAR foci to untreated levels **(Figure 2B)**, suggesting that most PARylation events induced by 5hmdC associated to ongoing replication are dependent on SMUG1. Consistent with the loss of PARylation associated to replication, loss of SMUG1 also abolished 5hmdC-induced heightened SSBs observed in *Fancd2^-/-^*cells, detected by alkaline comet assay **(Figure 2C),** suggesting that 5hmdC-induced ssDNA gap formation is dependent on SMUG1. Moreover, SMUG1 depletion restored replication fork progression and cell viability in 5hmdC-treated *Fancd2^-/-^* cells to *wild type* levels **(Figure 2D and 2E)**, demonstrating that SMUG1 loss suppresses ssDNA gap accumulation, replication fork instability and cell lethality observed in 5hmdC-treated HRD cells, most likely throughout abolishment of AP site formation.

**Figure 2.**
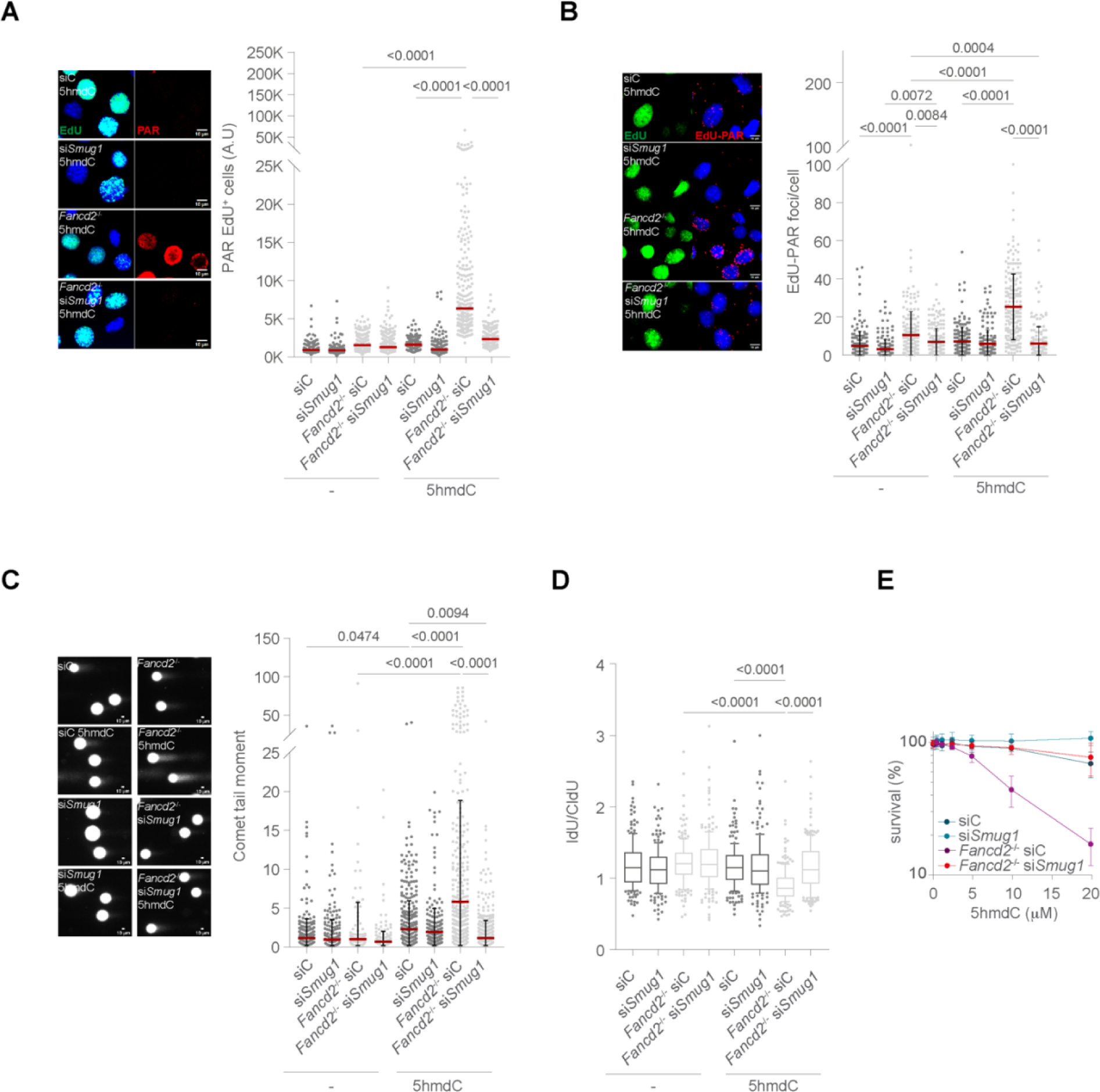
5hmdC-mediated HRD genomic instability is dependent on SMUG1. A. *Left*, representative PAR (red) immunofluorescence images of EdU^+^ (green) *wild type*, *Fancd2^- /-^*, si*Smug1* and *Fancd2^-/-^* si*Smug1* cells exposed to 5hmdC (10µM) for 3h. DAPI (blue) stains nuclear DNA. *Right*, plot depicting PAR mean intensity signal per nucleus. B. *Left*, representative images of EdU^+^-PAR foci (red) by SIRF assay from *wild type*, *Fancd2^-/-^*, si*Smug1* and *Fancd2^-/-^*si*Smug1* cells exposed to 5hmdC (10µM) for 3h. DAPI (blue) stains nuclear DNA and EdU (green) stains S-phase cells. *Right*, plot depicting EdU-PAR foci per nucleus (n=2. C. *Left*, representative images of alkaline comet assay from *wild type*, *Fancd2^-/-^*, si*Smug1* and *Fancd2^-/-^* si*Smug1* cells following 5hmdC treatment (10µM) for 3h. *Right*, plot depicting comet tail moment per cell (n=2. D. Box plot representing the frequency of IdU/CldU ratio of *wild type*, *Fancd2^-/-^*, si*Smug1* and *Fancd2^-/-^* si*Smug1* cells after 5hmdC treatment (40µM) for 30 min (n=200 of each 3 biological replicates,. E. MTT cell proliferation assay of *wild type*, *Fancd2^-/-^*, si*Smug1* and *Fancd2^-/-^* si*Smug1* cells, exposed to the indicated dose of 5hmdC for 3 days (n=4, mean ± s.d.).

Mammalian cells contain two class-II AP endonucleases APE1 and APE2(*60, 61*). While APE1 possess strong AP endonuclease but weak 3’-phosphodiesterase and 3’-5’exonuclease activities, APE2 exhibits strong *in vitro* 3’-phosphodiesterase and 3’-5’exonuclease activities, stimulated *in vivo* by PCNA binding, but weak AP endonuclease (*61*). We firstly examined the contribution of APE1 knockdown hardly had any effects on nuclear PARylation of untreated cells, but 5hmdC- dependent nuclear PARylation in EdU^+^ *Fancd2^-/-^* cells significantly decreased to near *wild type* levels **(Figure 3A and supplementary figure 2C)**. Moreover, SIRF assay showed a major role of APE1 in mediating the PARylation events associated to DNA synthesis. APE1 loss significantly decreased the spontaneous EdU-PAR foci in *Fancd2^-/-^*cells, suggesting that APE1 is responsible for a proportion of spontaneous ssDNA gaps observed in HRD cells. Moreover, APE1 depletion significantly decreased EdU-PAR foci to *wild type* levels in both 5hmdC-treated cell lines **(Figure 3B)**, suggesting that APE1 is responsible for most, if not all, 5hmdC-induced EdU-PAR foci. In agreement with these results, alkaline comet assay showed that APE1 knockdown significantly reduced 5hmdC-induced SSBs observed in *Fancd2^-/-^*cells to almost untreated levels **(Figure 3C)**, suggesting that APE1 is the main endonuclease responsible for most 5hmdC-induced SSBs in *Fancd2^-/-^* and, to a lesser extent, in *wild type* cells. 5hmdC-mediated replication fork impairment and cell lethality phenotypes observed in *Fancd2^-/-^* cells were both substantially suppressed by APE1 loss **(Figure 3D and 3E)**, suggesting that APE1-dependent SSBs account for the replication fork stalling and viability loss phenotypes observed in 5hmdC-treated *Fancd2*^-/-^ cells. As APE1 loss does not fully restore cell viability nor fork dynamics, these results suggest that APE1 may play additional roles during stability or resumption of replication forks. As APE2 loss has been shown to provide synthetic lethal phenotype in HRD deficient cells (*62, 63*), we decided to determine PAR levels in total or EdU^+^ APE2 or APE1 APE2 co-depleted *Fancd2*^-/-^ cell populations. Similar to APE1 depletion, APE2 depletion also reduced 5hmdC-mediated nuclear PARylation in total or S- phase (EdU^+^) *Fancd2^-/-^* cells populations **(supplementary figure 4A and 4B)**, but to a lesser extent than APE1 loss, and APE1 and APE2 codepletion showed an epistatic relationship, as PAR levels in APE1 APE2 co-depleted S-phase cells were similar to those observed in APE1-depleted *Fancd2*^-/-^ cells. Moreover, APE1 or APE2 knockdown completely suppressed 5hmdC-induced EdU-PAR foci and SSBs to untreated levels **(Figure 3F and supplementary figure 4C)**, suggesting that APE1, in combination with APE2, are responsible for the increased PARylation and SSBs associated to nascent strand during 5hmdU removal. Consistent with this notion, APE1 and APE2 showed an epistatic relationship not only on 5hmdC-mediated PAR levels in EdU^+^ *Fancd2^-/-^* cells but also in the suppression of 5hmdC-mediated lethality **(Figure 3E)**, suggesting that APE1 and APE2 function together to process 5hmdU-derived AP sites, likely generating ssDNA gaps.

**Figure 3.**
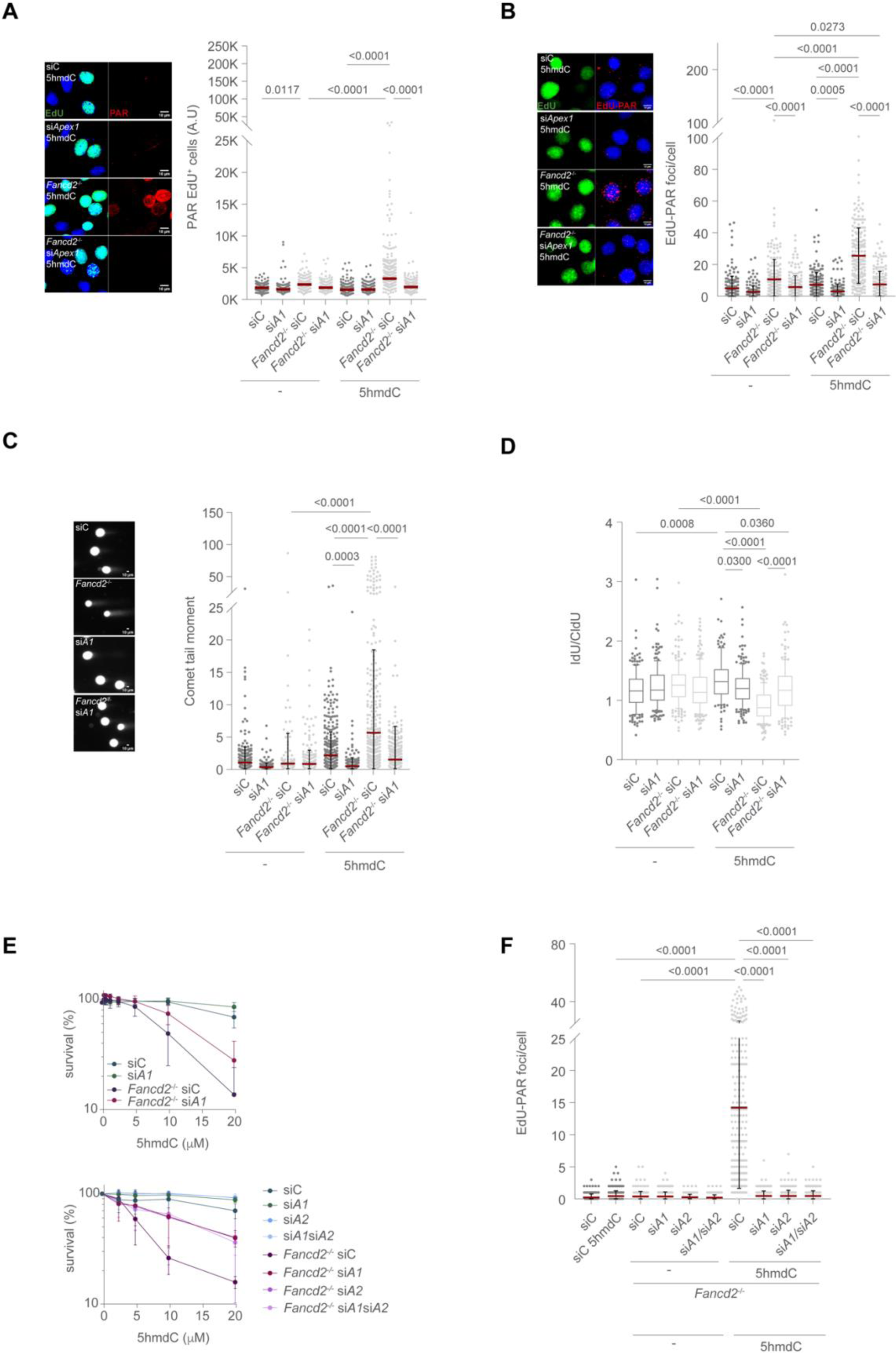
APEX1 and APEX2 contribute to 5hmdC-induced ssDNA gap generation in HRD A. *Left*, representative PAR (red) immunofluorescence images of APE1-depleted *wild type* or *Fancd2^-/-^* cells exposed to 5hmdC (10µM) for 3h. DAPI (blue) stains nuclear DNA and EdU (green) stains S-phase cells. *Right*, plot depicting PAR mean intensity signal per EdU^+^ nucleus. B. *Left*, representative images of EdU-PAR foci (red) by SIRF assay from *wild type*, *Fancd2^-/-^*, si*Apex1* or *Fancd2^-/-^*si*Apex1* cells exposed to 5hmdC (10µM) for 3h. DAPI (blue) stains nuclear DNA and EdU (green) stains S-phase cells. *Right*, plot depicting EdU-PAR foci per nucleus. C. *Left*, representative images of alkaline comet assay of APE1 (si*A1*) knocked down *wild type* or *Fancd2^-/-^* cells exposed to 5hmdC (10µM) for 3h. *Right*, plot depicting comet tail moment per cell. D. Box plot of the frequency of IdU/CldU ratio of APE1-depleted *wild type* or *Fancd2^-/-^* cells upon 5hmdC treatment (40µM) for 30 min (n=200 of each 2 biological replicates. E. *Top*, MTT cell proliferation assay of APE1-depleted *wild type* or *Fancd2^-/-^* cells exposed to the indicated dose of 5hmdC for 3 days (n=9, mean ± s.d). *Bottom*, MTT cell proliferation assay of APE1- or APE2-depleted *wild type* or *Fancd2^-/-^* cells exposed to the indicated dose of 5hmdC for 3 days. F. Plot depicting EdU-PAR foci per nucleus by SIRF assay from *wild type*, *Fancd2^-/-^*, si*Apex1*, si*Apex2*, *Fancd2*^-/-^ si*Apex1* and *Fancd2^-/-^* si*Apex2* cells exposed to 5hmdC (10µM) for 3h (n=2).

### 5hmdU-derived ssDNA gap persistence at replication forks exacerbate genomic instability and lethality in HRD cells

Inefficient ligation of nascent OF by the XRCC1/POLβ/LIG1 preligation complex in unperturbed S-phase triggers ssDNA gap persistence and PARP1-dependent PARylation events, which requires XRCC1-mediated PARP1 removal(*64*). In a similar manner, APE1-mediated incision of the AP sites in the context of replication forks may mimic persistent ssDNA gap formation. Moreover, APE2-mediated ssDNA gap extension may enhance ATR-dependent checkpoint signalling to promote HR(*65, 66*). We therefore reasoned that defective ssDNA gap processing either by excessive PARP1 retention upon Olaparib treatment, or by loss of XRCC1 or PARP1, would exacerbate ssDNA gap persistence and cell lethality in our settings. Indeed, XRCC1 knockdown further increased nuclear PARylation in 5hmdC-treated total or EdU^+^ *Fancd2^-/-^* cell populations whilst remained unaffected in *wild type* cells **(Figure 4A and supplementary figure 2D).** Similarly, SIRF assay showed exacerbation of EdU-PAR foci in XRCC1-knockdown *Fancd2*^-/-^treated cells **(Figure 4B)**, suggesting that ssDNA gaps associated to replication forks in the absence of FANCD2 persist longer or accumulate at larger extent upon XRCC1 loss. In agreement with this notion, either PARP1 or XRCC1 depletion, or PARP1 trapping by Olaparib(*15*), which result in inefficient repair of ssDNA gaps, further exacerbated 5hmdC-mediated *Fancd2^-/-^* cell lethality **(Figure 4C)**, without further impacting on 5hmdC-mediated *Fancd2^-/-^* replication fork stalling **(Figure 4D)**. As ssDNA gaps emerging from stalled replication forks by bulky base adducts stimulate post-replicative SCEs (*37*), we examined if 5hmdC-mediated ssDNA gaps impacted on SCEs in HRD cells. To detect SCEs derived from ssDNA gaps formed specifically in nascent DNA, cells were exposed to 5hmdC less than one cell cycle length (≈12 hours) before harvesting and SCEs were scored. We found that upon 5hmdC treatment, *Fancd2^-/-^*cells showed a significant increase in SCEs compared to untreated cells **(Figure 4E)**, suggesting that ssDNA gaps in nascent strands in the current cell cycle are a source of SCEs in HRD cells. Moreover, knockdown of PARP1 or XRCC1 increased 5hmdC-induced SCEs whilst SMUG1 or APE1 depletion blunted this response **(Figure 4E)**. These results suggest that defective repair of persistent replicative 5hmdC- mediated ssDNA gaps account for the increased replication fork-associated PARylation, SSBs, SCEs and cytotoxicity observed in *Fancd2^-/-^* cells.

**Figure 4:**
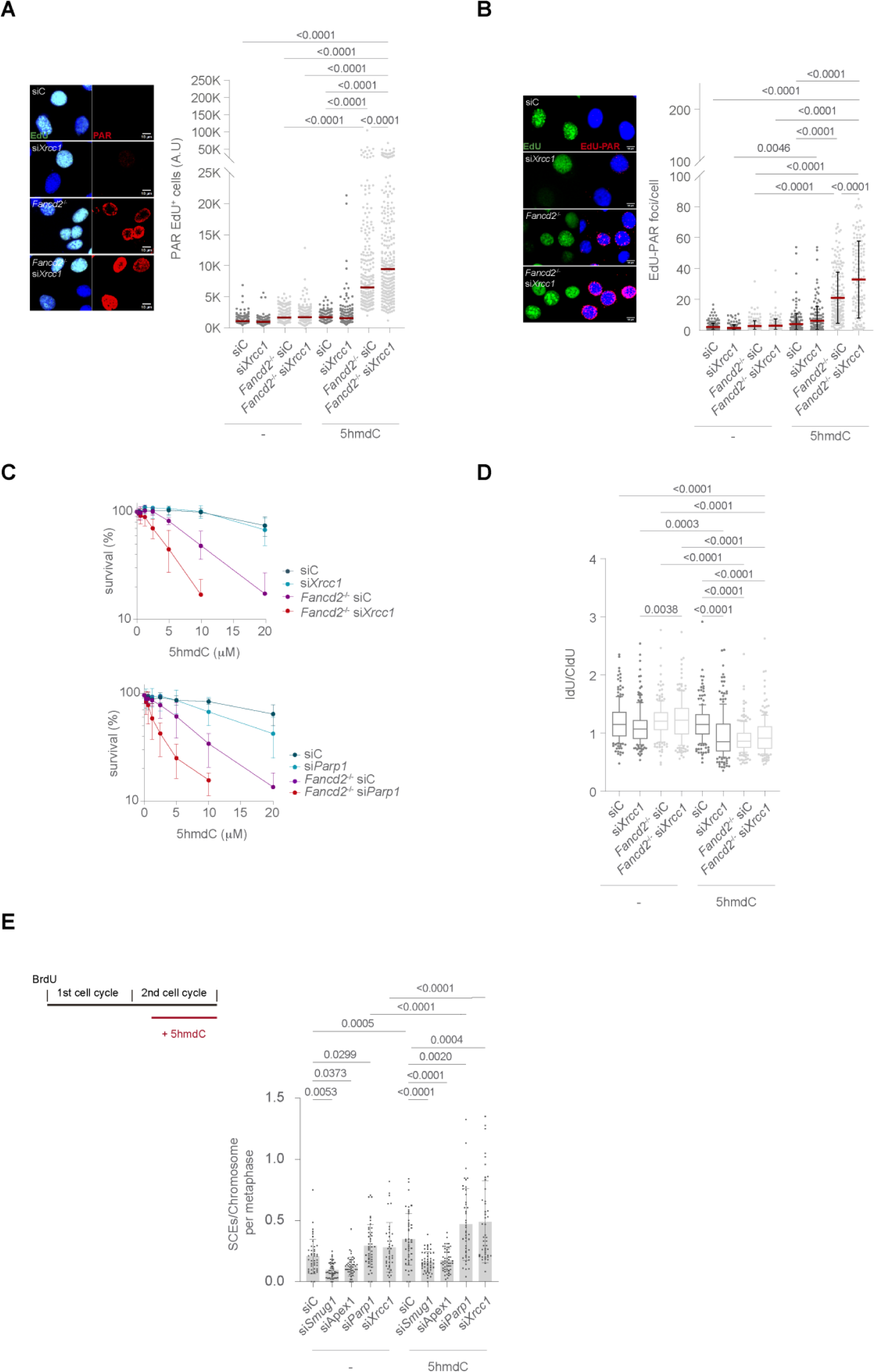
XRCC1 depletion exacerbate 5hmdC-induced genomic instability and lethality of HRD cells. A. *Left*, representative PAR (red) immunofluorescence images of XRCC1-depleted *wild type* or *Fancd2^-/-^* cells exposed to 5hmdC (10µM) for 3h. DAPI (blue) stains nuclear DNA and EdU (green) stains S-phase cells. *Right*, plot depicting PAR mean intensity signal per nucleus. B. *Left*, representative images of EdU-PAR foci (red) by SIRF assay of XRCC1-depleted *wild type* or *Fancd2^-/-^* cells exposed to 5hmdC (10µM) for 3h. DAPI (blue) stains nuclear DNA and EdU (green) stains S-phase cells. *Right*, plot depicting EdU-PAR foci per nucleus. C. *Top*, MTT cell proliferation assay of *wild type* or *Fancd2^-/-^* cells lacking XRCC1 exposed to the indicated dose of 5hmdC for 3 days (n=9, mean ± s.d.). *Bottom*, MTT cell proliferation assay of *wild type*, *Fancd2^-/-^*, si*Parp1*, and *Fancd2^-/-^* si*Parp1* cells, exposed to the indicated dose of 5hmdC for 3 days (n=5, mean ± s.d.). D. Box plot of the frequency of IdU/CldU ratio of XRCC1-depleted *wild type* or *Fancd2^-/-^* cells after 5hmdC treatment (40µM) for 30 min (n=200 of each 2 biological replicates. E. *Top left*, scheme of the SCE assay. *Right*, bar plot of SCE/chromosome per metaphase from SMUG1-, APEX1-, XRCC1- or PARP1-depleted *Fancd2^-/-^* cells after exposure to 5hmdC (10µM) for 12h (n=50 of each of 2 biological replicates; bar represents mean ± s.d.).

### HMCES-DPCs are responsible for 5hmdC-mediated ssDNA gaps in HRD cells

Presence of uracil, either by misincorporation on nascent DNA strand or by APOBEC-mediated 5dC deamination on template strand, is recognized and removed by UNG glycosylases during BER, leaving AP sites and other intermediate DNA ends that hamper replication fork progression(*48, 49, 53, 67*). HMCES covalently reacts to aldehydic form of AP sites to regulate APE1-mediated endonucleolytic cleavage, which avoids cytotoxic DSB formation at the replication fork whilst promoting DDT base lesion bypass mechanisms such as TLS(*50, 51, 53*). We therefore reasoned that HMCES would react to 5hmdU-derived AP sites arising during 5hmdC treatment. In agreement with this notion, in untreated conditions we observed a higher proportion of Flag-tagged HMCES positive nuclei in *Fancd2*^-/-^ cells than in *wild type* cells **(Figure 5A)**. Moreover, *Fancd2^-/-^* cells displayed a brighter nuclear HMCES signal compared to its *wild type* counterpart under spontaneous, and methyl methanesulfonate (MMS) or 5hmdC exposure, chemicals known to induce AP site formation. Moreover, chromatin-bound HMCES levels directly correlated with 5hmdC misincorporation on nascent strand in a dose-dependent manner, reaching higher levels in *Fancd2*^-/-^ than in *wild type* cells **(Figure 5B)**. These data suggest that *Fancd2*^-/-^ cells present higher endogenous levels of HMCES-bound AP sites than *wild type* cells, which increase upon MMS or 5hmdC treatments in a dose-dependent manner. To confirm these data, we generated cell lines overexpressing either wild type (WT), Cys2Ala (C2A) or Arg212Glu (R212E) Flag-tagged HMCES constructs and examined their chromatin retention activity upon MMS or 5hmdC exposure. In agreement with their reported *in vitro* biochemical activities(*50*), compared to HMCES -WT, HMCES-C2A showed defective chromatin recruitment upon 5hmdC or MMS exposure while HMCES-R212E completely abolished chromatin binding **(Figure 5C)**. These data suggest that AP site on nascent DNA strand from 5hmdU processing by BER is recognized by HMCES, which forms a HMCES-DPC. We next reasoned that loss of HMCES-DPC would increase the APE1- mediated AP sites conversion to cytotoxic ssDNA gaps on nascent DNA strand right at or behind the fork, resulting in a synthetic lethal phenotype in HRD cells. To our surprise, HMCES knockdown dramatically decreased 5hmdC-induced nuclear PARylation in *Fancd2^-/-^* total or EdU^+^ cell populations **(supplementary figure 5A and figure 5D)**, possibly due to a role of HMCES outside replication forks. EdU-PAR SIRF assay confirmed the absence of PARylation in HMCES-depleted *Fancd2^-/-^* S-phase cells **(Figure 5E)**, suggesting that HMCES-DPC on nascent DNA strand is responsible for heightened PAR levels associated to replication forks in HRD cells. Consistent with the protective role of HMCES during replication (*50*), nuclear γ-H2AX levels increased upon HMCES depletion, which was further exacerbated in *Fancd2^-/-^* cells **(supplementary figure 5B)**. Nevertheless, γ-H2AX levels remained relatively unchanged during 5hmdC treatment, suggesting that conversion of 5hmdU-derived AP site to ssDNA gaps in nascent DNA upon HMCES depletion do not result in cumulative DSBs **(supplementary figure 5B)**. The contribution of HMCES loss to nascent ssDNA gap formation was also examined by alkaline comet assay. HMCES knockdown had barely any effect on the frequency of SSBs in either treated or untreated *wild type* cells. However, 5hmdC-induced SSBs observed in *Fancd2^-/-^ s*i*Hmces* cells significantly decreased to *wild type s*i*Hmces* treated levels, not reaching *Fancd2^-/-^* siC untreated levels **(Figure 5F)**, suggesting that HMCES-DPCs are responsible for a large subset of 5hmdC-induced ssDNA gaps observed in *Fancd2^-/-^* cells.

**Figure 5.**
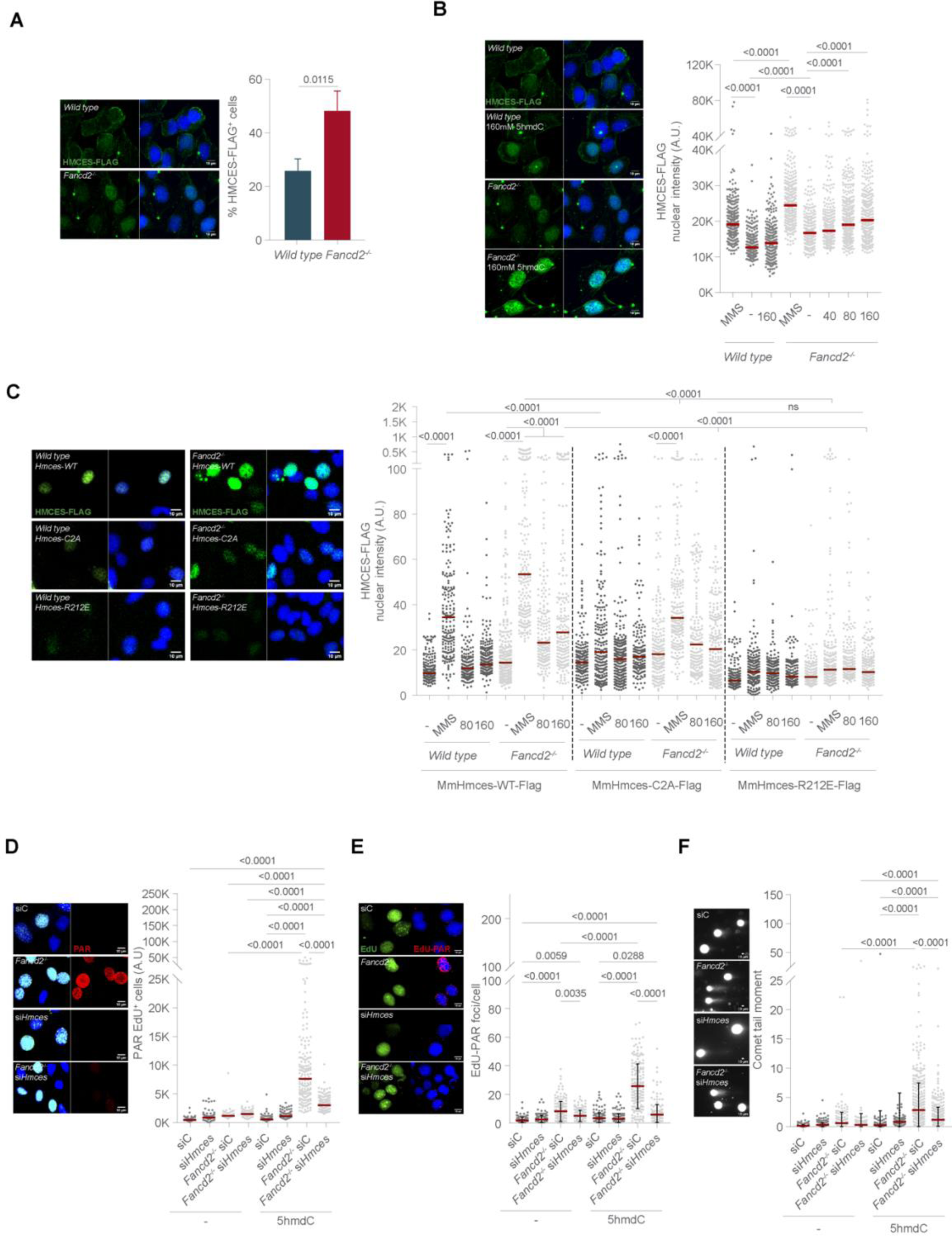
HMCES accumulates in nuclei and is responsible for 5hmdC-mediated genomic instability observed in HRD cells A. *Left*, representative immunofluorescence images of Flag-tagged HMCES (green) positive nuclei from *wild type* or *Fancd2*^-/-^ cells. *Right*, percentage of HMCES-FLAG^+^ *wild type* and *Fancd2*^-/-^ cells. B. *Left*, representative immunofluorescence images of HMCES positive nuclei from *wild type* or *Fancd2*^-/-^ cells upon 5hmdC (160µM) plus MG-132 (10µM) combined treatments for 3h. *Right*, HMCES-FLAG mean intensity signal per nucleus from *wild type* and *Fancd2*^-/-^ cells upon 5hmdC (indicated doses for 3h) or MMS (0.5mM, 30min). Treatments were combined with MG-132 (10µM). C. *Left*, images of FLAG-tagged HMCES positive nuclei from *wild type* or *Fancd2*^-/-^ cells, expressing MmHMCES-WT-FLAG, MmHMCES-C2A-FLAG or MmHMCES-R212E-FLAG upon 5hmdC exposure (160µM, 3h). *Right*, plot depicting HMCES-FLAG mean intensity signal per nucleus from *wild type* or *Fancd2*^-/-^ cells expressing MmHMCES-WT-FLAG, MmHMCES- C2A-FLAG or MmHMCES-R212E-FLAG upon 5hmdC (indicated doses for 3h) or MMS (0.5mM). Treatments were combined with MG-132 (10µM). D. *Left*, representative PAR (red) immunofluorescence images of HMCES-depleted *wild type* or *Fancd2^-/-^*cells exposed to 5hmdC (10µM) for 3h. DAPI (blue) stains nuclear DNA. EdU (green) stains S-phase cells. *Right*, PAR mean intensity signal per nucleus. E. *Left*, representative images of EdU-PAR foci by SIRF assay from HMCES-depleted *wild type* or *Fancd2^-/-^* cells exposed to 5hmdC (10µM) for 3h. DAPI (blue) stains nuclear DNA. EdU (green) stains S-phase cells. *Right*, plot depicting EdU-PAR foci per nucleus. F. *Left*, representative images of alkaline comet assay from HMCES-depleted *wild type* or *Fancd2^-/-^* cells following 5hmdC treatment (10µM) for 3h. *Right*, plot depicting comet tail moment per cell.

Next, we examined the impact of HMCES loss in 5hmdC-mediated perturbed replication fork dynamics and chromosomal aberrations observed in *Fancd2^-/-^* cells. As previously reported, 5hmdC induced replication fork impairment in HRD cells(*15*). Nevertheless, loss of HMCES restored *Fancd2^-/-^* replication fork progression and replication fork symmetry to *wild type* levels **(Figure 6A and 6B)**. Moreover, 5hmdC-induced heightened chromosomal aberrations were markedly reduced upon HMCES depletion **(Figure 6C)**, specifically those of radial chromosome, which arose at the expenses of chromosomal gaps **(supplementary figure 6A)**. Similarly, lethality of HRD cells induced by 5hmdC was also significantly suppressed upon HMCES depletion **(Figure 6D)**, indicating that 5hmdU-derived HMCES-DPCs formed during BER intermediates are responsible for replication fork defects, chromosomal instability and cell lethality in HRD cells. To study the suppression of ssDNA gaps by HMCES loss, we attempted to carry out the S1 nuclease DNA fiber assay under these conditions in MEFs but failed to obtain reproducible results. To overcome this, we successfully CRISPR-CAS9 knocked out FANCD2, HMCES or both genes in the human RPE- 1 cell line **(supplementary figure 7)** and carried out the S1 nuclease assay. Consistent with published data, HMCES loss did not impact on unperturbed replication fork progression (*51*) or in the amount of ssDNA gaps measured by the S1 nuclease assay, suggesting that unprotection of spontaneous AP sites upon HMCES loss do not affect overall levels of ssDNA gaps **(Figure 6E)**. Exposure to 5hmdC (10µM) significantly affected IdU track length in the absence of FANCD2, suggesting that FANCD2 is required to avoid ssDNA gap formation. However, and consistent with our previous data, HMCES depletion suppressed 5hmdC-induced ssDNA gap formation observed in *FANCD2*^-/-^ cells **(Figure 6E)**, suggesting that HMCES is responsible for the heightened 5hmdC- mediated ssDNA gaps observed in HRD cells. We also exploited breast tumour databases to find any correlation of HMCES loss and tumour progression. We found that low HMCES expression levels correlated with lower overall survival in a breast cancer cohort (TCGA=1089 pts) by RNA-seq or array **(Figure 6F and 6G)** supporting the notion that HMCES loss may impact in tumour aggressiveness and breast cancer patient outcome.

**Figure 6.**
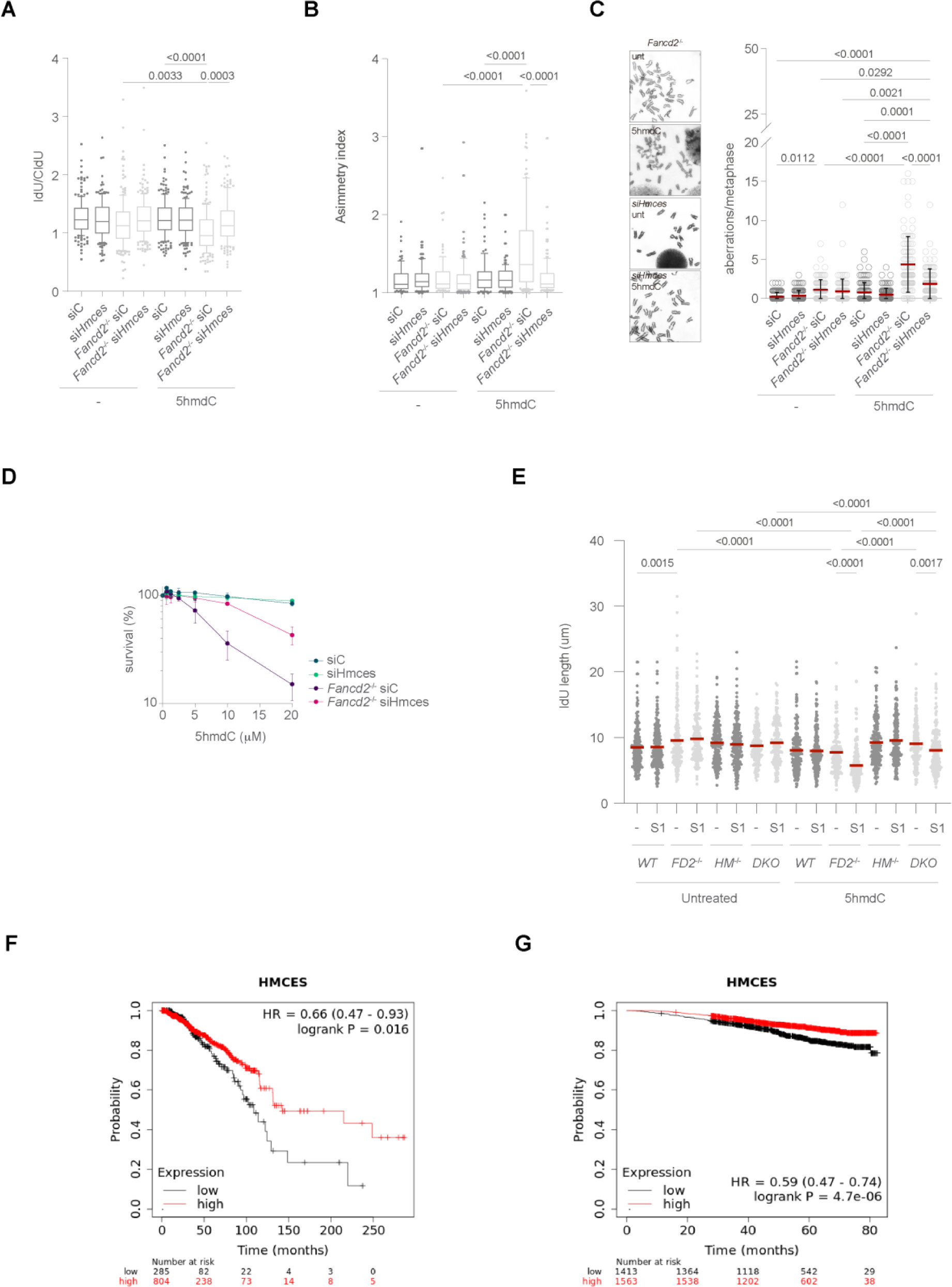
HMCES depletion rescues 5hmdC-induced replication fork defects and ssDNA gap accumulation in HRD cells. A. Box plot of the frequency of IdU/CldU ratios of HMCES-deficient *wild type* or *Fancd2^-/-^* cells upon 5hmdC treatment (40µM) for 30 min (n=200 of each 3 biological replicates). B. DNA fiber asymmetry index from HMCES-depleted *wild type* or *Fancd2^-/-^* cells upon 5hmdC treatment (40µM) for 30 min. C. *Left*, chromosome aberrations from HMCES-knockdown cells following 5hmdC treatment (10µM) for 40h. *Right*, plot of total chromosome aberrations/metaphase (n=50 of each of 2 biological replicates; bar represents mean ± s.d.). D. MTT cell proliferation assay of HMCES-depleted *wild type* or *Fancd2^-/-^*cells, exposed to the indicated dose of 5hmdC for 3 days (n=3, mean ± s.d.). D. Dot plot of IdU track length of *WT*, *FANCD2^-/-^* (*FD2^-/-^*), *HMCES^-/-^* (*HM^-/-^*) or *FANCD2^-/-^HMCES^-/-^* (*DKO*) cells upon 5hmdC treatment (10µM) for 30 min. S1 treatment was carried out for 30 min. (n=200 of each 2 biological replicates). E and F. Levels of HMCES mRNA and overall survival in a breast tumour cohort of the human TCGA repository

HMCES-DPC on template DNA strand has been described to show an epistatic relationship over APE1 incision to avoid toxic DSB formation(*50, 52*). To study the epistatic relationship of HMCES, APE1 and APE2 upon 5hmdC exposure, we examined S-phase nuclear PARylation and viability phenotypes upon depletion of HMCES, APE1, APE2 or their combinations. Nuclear PARylation of HMCES APE1 double knockdown EdU^+^ cells resembled the one seen in *Fancd2^-/-^* APE1 single knockdown cells **(Figure 7A)**, whereas PARylation observed in HMCES APE2 double knockdown *Fancd2^-/-^* EdU^+^ cells resembled the one observed in *Fancd2^-/-^* HMCES single knockdown EdU^+^ cells **(Figure 7B)**. Moreover, *Fancd2^-/-^*cell survival also showed an epistatic suppression upon knockdown of either HMCES, APE1 or APE2 **(Figure 7C and 7D)**, suggesting that HMCES, APE1 and APE2 function in the same genetic pathway during the processing of 5hmdU from nascent DNA.

**Figure 7.**
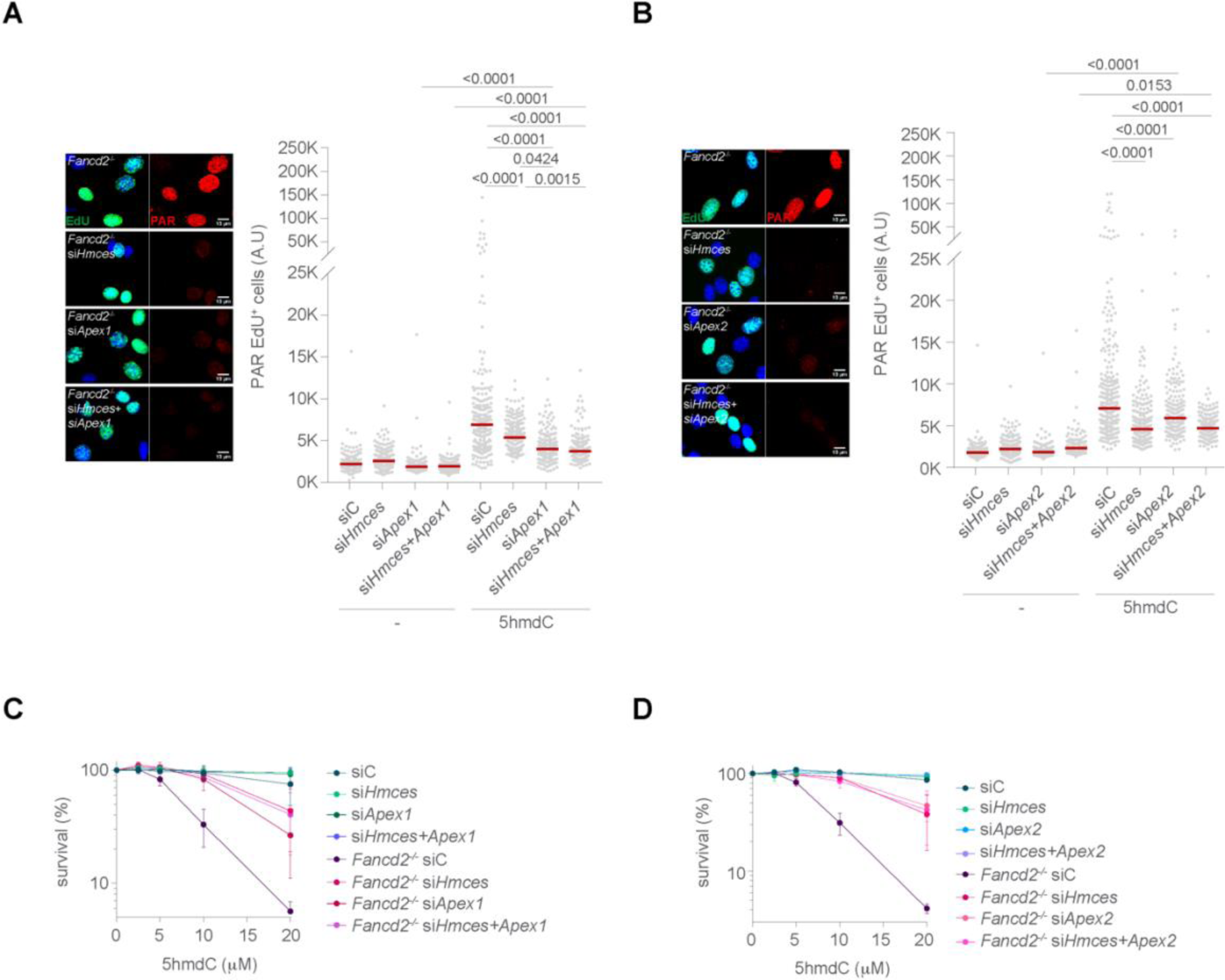
A. HMCES, APEX1 and APEX2 function in a common BER genetic pathway during 5hmdU removal A. *Left*, representative PAR (red) immunofluorescence images from *wild type* or *Fancd2^-/-^* cells upon depletion of HMCES, APEX1 or their combinations exposed to 5hmdC (10µM) for 3h. DAPI (blue) stains nuclear DNA and EdU (green) stains S-phase nuclei. *Right*, plot depicting PAR mean intensity signal per nucleus. B. *Left*, representative PAR (red) immunofluorescence images from *wild type* or *Fancd2^-/-^* cells upon depletion of HMCES, APEX2 or their combinations exposed to 5hmdC (10µM) for 3h. DAPI (blue) stains nuclear DNA and EdU (green) stains S-phase cells. *Right*, plot depicting PAR mean intensity signal per nucleus. C. MTT cell proliferation assay of HMCES-, APEX1- or HMCES APEX1-depleted *wild type* or *Fancd2^-/-^*cells, exposed to the indicated dose of 5hmdC for 3 days (n=5, mean ± s.d.). D. MTT cell proliferation assay of HMCES-, APEX2- or HMCES APEX2-depleted *wild type* or *Fancd2^-/-^* cells, exposed to the indicated dose of 5hmdC for 3 days.

### PRIMPOL mediates a subset of 5hmdC-mediated ssDNA gaps

It has previously been reported that replication fork stalling by bulky base adducts in mammalian cells generates post-replicative ssDNA gaps predominantly throughout the primase and polymerase activities of PRIMPOL(*37, 38*). These ssDNA gaps can be filled in either by the REV1-POLζ or POLQ TLS polymerases, or by alternative error-free recombination involving RAD51. We therefore examined the contribution of PRIMPOL to 5hmdC-dependent ssDNA gap formation observed in our cell lines. PRIMPOL depletion had little effect on PARylation levels in untreated EdU^+^ *Fancd2^-/-^* cells but significantly suppressed 5hmdC-mediated PARylation, not reaching *wild type* levels **(supplementary figure 8A)**. PRIMPOL depletion partially suppressed 5hmdC-induced EdU-PAR SIRF foci in *Fancd2^-/-^* cells **(Figure 8A)**, suggesting that a proportion of 5hmdC-induced ssDNA gaps depends on PRIMPOL. Moreover, comet assay of PRIMPOL-depleted cells showed a decreased in ssDNA gaps independently of the genotype under study. Whereas PRIMPOL depletion completely abolished 5hmdC-dependent SSBs in *wild type* cells, there was still a significant increase of SSBs in PRIMPOL-depleted *Fancd2*^-/-^ cells **(Figure 8B)**. This situation contrasted to APE1-depleted cells, whereby APE1 depletion completely abolished 5hmdC-induced ssDNA gaps in both backgrounds, indicating that, in addition to PRIMPOL-mediated ssDNA gaps, 5hmdC induces other types of ssDNA gaps, probably due to unrepaired APE1-mediated SSBs. Consistent with this data, PRIMPOL partially accounted for the 5hmdC-induced lethality observed in *Fancd2^-/-^* cells, and either APE1 PRIMPOL or HMCES PRIMPOL double knockdown cells displayed the same viability as PRIMPOL single depleted *Fancd2^-/-^* cells **(Figure 8C)**, suggesting that a subset of ssDNA gaps arising from HMCES-DPC toxic lesions are PRIMPOL-dependent whereas other gaps are independently formed. As PRIMPOL-dependent ssDNA gaps at nascent DNA strand induce SCEs upon exposure to bulky adducts, we examined the contribution of HMCES and PRIMPOL to SCEs induced by 5hmdC. PRIMPOL loss suppressed heightened SCEs induced by 5hmdC in *Fancd2^-/-^* cells. Strikingly, HMCES loss also blunted 5hmdC-dependent SCEs in *Fancd2^-/-^* cells, and co-depletion of HMCES and PRIMPOL showed an epistatic phenotype **(Figure 8D)**, suggesting that nascent strand ssDNA gaps arising from 5hmdC-induced HMCES- DPCs formed during the ongoing cell cycle are responsible for PRIMPOL-mediated heightened SCEs and suggest a more complex role of HMCES at protecting AP sites than previously anticipated.

**Figure 8.**
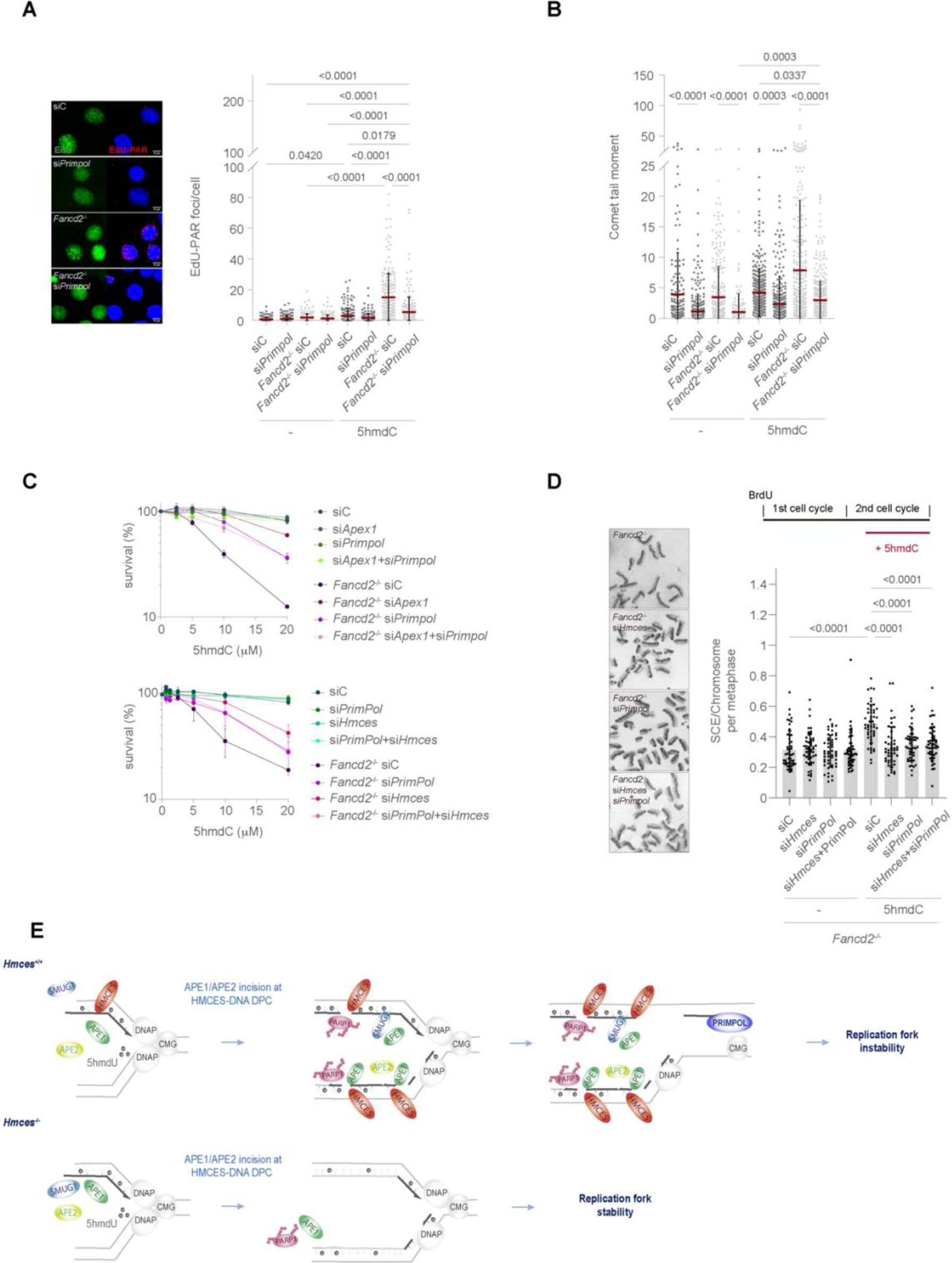
HMCES is responsible for PRIMPOL-dependent ssDNA gap accumulation. A. *Left*, representative PAR (red) immunofluorescence images of *wild type*, *Fancd2^-/-^*, si*Primpol* and *Fancd2^-/-^* si*Primpol* cells exposed to 5hmdC (10µM) for 3h. DAPI (blue) stains nuclear DNA. EdU (green) stains S-phase cells. *Right*, plot depicting PAR mean intensity signal per nucleus. B. *Left*, representative images of EdU-PAR foci (red) by SIRF assay from *wild type*, *Fancd2^-/-^*, si*Primpol* and *Fancd2^-/-^* si*Primpol* cells exposed to 5hmdC (10µM) for 3h. DAPI (blue) stains nuclear DNA. EdU (green) stains S-phase cells. *Right*, plot depicting EdU-PAR foci per nucleus. C. *Top*, MTT cell proliferation assay of *wild type*, *Fancd2^-/-^*, si*Ape1*, si*Primpol*, *Fancd2^-/-^* si*Ape1, Fancd2^-/-^* si*Primpol* and *Fancd2^-/-^* si*Primpol* si*Ape1* cells exposed to the indicated dose of 5hmdC for 3 days. *Bottom*, MTT cell proliferation assay of *wild type*, si*Hmces*, si*Primpol*, si*Hmces Primpol*, *Fancd2^-/-^*, *Fancd2^-/-^* si*Hmces*, *Fancd2^-/-^* si*Primpol* and *Fancd2^-/-^* si*Hmces Primpol* cells exposed to the indicated dose of 5hmdC for 3 days. D. *Left*, representative images of SCEs from *Fancd2^-/-^*, *Fancd2^-/-^* si*Hmces*, *Fancd2^-/-^* si*Primpol* and *Fancd2^-/-^* si*Hmces Primpol* cells exposed to 5hmdC (10µM) for 12h. *Top right*, scheme of SCE assay. *Bottom right*, bar plot of SCE/chromosome per metaphase (n=50 of each of 2 biological replicates; bar represents mean ± s.d.). E. Working model depicting the mechanism of replication fork instability by HMCES during removal of 5hmdU in the absence of the replication fork maintenance FA pathway pathway.

## Discussion

Chemical or physical modifications of DNA bases in addition to uracil (and its derivative forms) misincorporation are perhaps the major source of DNA lesions in cells. These harmful DNA lesions appear at estimated frequency of 10-50 x 10^4^ cell^-1^ day^-1^ (*25*). Due to their intrinsic chemical instability and to their deleterious consequences, AP sites are rapidly converted to SSBs by either the AP lyase activity or by the AP endonuclease activity of certain class of nucleases. Therefore, the repair of base alterations, AP sites and SSBs is perhaps the most important repair process responding to oxidative and alkylating base damages (*68, 69*), and the molecular basis of BER has been a long-standing question in the genome stability field. More intriguingly, is how BER repaired these damaged bases in the context of the replication forks, as AP sites and SSBs are potent replication fork blocking lesions, which when left unrepaired, hinder replication fork progression and stability. For these reasons, the study of the molecular mechanisms controlling ssDNA gap formation is of paramount interest in the context of HR-deficiency as many of the factors involved in HR (BRCA1, BRCA2, RAD51, FANCD2…) are also important for maintaining replication fork stability, as well as the use of drugs trapping PARPs in ssDNA gaps behind replication forks, which are clinically relevant to eradicate HRD tumor cells.

Recent work has identified the molecular mechanisms of dU, 5CldU or 5hmdU genotoxicity (*15, 44, 45, 48, 49*). During dU, 5CldU or 5hmdU misincorporation, the N-glycosylase activity of UNG1/2 or SMUG1 mediates efficient dU, 5CldU or 5hmdU base excision. UNG or SMUG1 N-glycosylase activities on dU or 5hmdU respectively generate AP sites, which when misrepaired, trigger replication fork conflicts due to persistent BER intermediates in template DNA. In the absence of AP site generation, unprocessed dU triggers an ATR-dependent response characterized by replication fork slowing and PRIMPOL-dependent ssDNA gap formation, presumable as a consequence of replication fork stalling (*49*). Our study complements the previous ones by reporting that HMCES-DPCs, generated upon 5hmdU misincorporation on the nascent strand, is an additional toxic BER intermediate challenging replication fork stability. By replication fork dynamics, detection of BER-mediated PARylation events associated to newly incorporated EdU, alkaline comet and viability assays, in combination with our genetic epistasis analysis, we conclude that HMCES-DPCs on nascent DNA promote incisions by APE1/APE2, which are responsible for stalling replication forks and the accumulation of postreplicative ssDNA gaps. We identify two sources of ssDNA gaps. One source comes from direct incisions at the HMCES-DPC by APE1/APE2, which probably accounts for ssDNA gaps at or behind the fork. The second source of ssDNA gaps is dependent on both APE1 and PRIMPOL activities and probably accounts for ssDNA gaps ahead the stalled fork, due to PRIMPOL-mediated repriming of DNA synthesis to accomplish genome duplication. *In vitro* assays have determined that HMCES binds to UDG-mediated AP sites in ssDNA to inhibit APE1 incision, while the self-reversal activity releases HCMES from dsDNA (*50, 55, 56*). Considering that *Fancd2*^-/-^ cells displayed higher chromatin retention of HMCES than *wild type* cells, which is dependent on both DNA binding and crosslinking activities of HMCES in accordance to the reported biochemical *in vitro* activities of HMCES (*50*), this data suggests that, in the absence of the FA/BRCA pathway, cells accumulate stretches of ssDNA with embedded AP-sites, which are protected by HMCES. Alternatively, nucleofilament formation by FANCD2 and its partner FANCI, might somehow protect genome integrity from AP sites accumulation to avoid toxic reactivity of AP sites or APE1-mediated SSBs.

In addition to the recently reported self-reversal activity of HMCES, proposed to occur on dsDNA, HMCES may also undergo FANCJ-mediated denaturation and subsequent proteasome or SPRTN-mediated proteolysis(*50, 54*). This replication fork-associated HMCES degradation is predicted to leave a peptide bound to the ribose through a hard-to-degrade thiazolidine bond. In this situation, APE1 has been reported to efficiently incise 5’to the DNA-peptide-crosslink. APE1 has been reported to display a prominent endonuclease over exonuclease activity, whilst APE2 shows an opposite behavior. Moreover, APE1 physically interact with APE2, PCNA and ATR, being the latter independently of RPA and promote checkpoint activation, and APE1 efficiently incises DNA- HMCES peptide crosslink(*65, 70–72*). In agreement with this, our data showed that APE1, in collaboration with APE2, mediate endonucleolytic incisions upon 5hmdC exposure. Our data shows that loss of APE1 suppressed the *Fancd2*^-/-^ 5hmdC-mediated EdU-PAR foci formation, SSBs accumulation by alkaline comet assay and heightened SCEs, to a similar extent as APE2 depletion, which displayed an epistatic behavior to APE1. The fact that APE2 depletion suppressed 5hmdC- induced SSBs in the alkaline comet assay to levels observed in APE1 knockdown cells strongly suggest that APE1 and APE2 activities are genetically linked. We favor a model in which AP sites arising from SMUG1-dependent activity at newly synthesized DNA would be protected by HMCES forming a HMCES-DPCs in both leading and lagging strands. Due to its strong *in vitro* ssDNA binding activity, we reasoned that HMCES would react to AP sites containing nascent ssDNA allowing HMCES-DPC formation at the fork. Once HMCES is partially degraded, APE1 through its phosphodiesterase activity, probably would incise 5’at the lesion site leaving a 5’-phosphate bound to ring-opened AP reacted to cys2-peptide through a thiazolidine bond. As a result, persistent 5’-peptide DNA or futile SSB ligation resembled the recently described replication-associated PARP1-mediated signaling of unligated Okazaki fragments, which would hinder replication fork progression leading to lethality of HRD cells.

HMCES has also been reported to play a key role in long-overhangs alternative-mediated end joining (Alt-EJ) pathway during class switch recombination in mice, a function requiring the ssDNA binding but not the crosslinking activities of HMCES(*73*). Alt-EJ is probably the ultimate repair mechanism to fix persistent DSBs in mitosis (*71, 74*), and is composed of end resection factors, POLQ, and BER sealing enzymes LIG1 and LIG3. Due to the fact that cells carrying a dysfunctional FA/BRCA pathway accumulate DSBs, these cells are reliant on mitotic Alt-EJ, whose deficiency caused by mutations in POLQ, LIG3 or ERCC1 efficiently induce HR-deficient cell death (*16, 18, 75*). Whereas low HMCES levels would decrease survival of HRD tumor cells, our data analysis using a large breast tumor cohort of the human TCGA repositorie, where HRD tumors are overrepresented, showed a strong direct correlation between overall survival and HMCES mRNA levels, suggesting that loss of HMCES may contribute to disease progression or relapse, which could be used as a marker of cancer susceptibility.

As a consequence of replication fork stalling by 5hmdU misincorporation, is the activation of replication fork rescue DDT mechanisms such as RFR, TS and TLS in HRD cells to promote replication completion (*33, 38, 45, 47*). Whereas RFR have been demonstrated to promote fork degradation in HRD cells (*39, 59, 76*), TS and PRIMPOL-mediated repriming are perhaps the chosen mechanisms to accomplished faithful DNA duplication whilst leaving cytotoxic ssDNA gaps behind replication forks (*47*). This ssDNA gaps can be subsequently filled in either a POLQ or REV1,3,7-dependent fashion (*38, 40*), or alternatively cleaved by endonucleases such as MRE11 to promote TS (*39*). Our data points out a role for PRIMPOL in a subset of 5hmdU-mediated ssDNA gaps. As previously observed by Piberger et al. (*37*), PRIMPOL-dependent ssDNA gap accumulation induced by large-adducted bases are mainly repair throughout a TS mechanism, leading to increased SCEs. In agreement with their observations, 5hmdC-induced HCMES-DPCs in nascent DNA also increased SCEs in HR-deficient *Fancd2*^-/-^ cells. Heightened 5hmdC-mediated SCEs observed in *Fancd2*^-/-^ cells were completely dependent not only on the PRIMPOL but also on HMCES, showing an epistatic relationship, suggesting that most SCEs in HRD-deficient cells arise from PRIMPOL-dependent ssDNA gaps. Moreover, depletion of SMUG1 or APE1, which generated AP sites or incisions at HMCES-DPCs suppressed 5hmdC-mediated SCEs, whereas loss of XRCC1 or PARP1 resulted in nascent strand ssDNA gap persistence exacerbating SCEs. Together, these results suggest that HMCES-DPCs formed in nascent DNA during 5hmdU removal induces PRIMPOL-dependent ssDNA gap formation and SCEs. As *Fancd2*^-/-^ cells are deficient in RAD51-mediated HR, it would be interesting to identify if RAD51, RAD52 or other paralogues promote strand invasion and faithful SCEs completion in these cells.

One of the limitations of our study is the lack of direct measurements of ssDNA in MEFs. After several attempts on setting up the S1 nuclease DNA fiber assay in MEFs, we could not obtain reproducible results. To overcome this issue, we generated RPE-1 cell lines knocked out for *HMCES*, *FANCD2* or both genes (*FANCD2 HMCES* DKO). S1 fiber assays reproducibly showed that 5hmdC treatment caused a significant accumulation of ssDNA gaps in *FANCD2*^-/-^ cells compared to *wild type* cells. In this cell type, HMCES loss largely suppressed 5hmdC-induced ssDNA gap accumulation upon S1 treatment, in agreement with our previous results in MEFs. However, our S1 nuclease fiber assay conditions did not allow us to discriminate the two types of ssDNA gaps. One possible explanation is that most ssDNA gaps detected by S1 nuclease assay are the ones generated by PRIMPOL activity, which are long enough as to accommodate the enzymatic activity, whereas the APE1-mediated SSBs are probably shorter. APE1-dependent ssDNA gap at 5’-DPC further extended by the robust 3’-5’exonuclease activity of APE2, might provide ssDNA substrate long enough for PARP signaling, but might require further extension by long-range resection nucleases (DNA2 or EXO1) to be accessible for S1 nucleases, as recently suggested by other labs (*77, 78*). This study highlights the diverse nature of ssDNA gap formation in nascent DNA and highlights endogenous HMCES-DPCs as novel source contributing to genomic instability with consequences beyond our expectations. Further work is needed to delineate their impact in tumorgenesis.

## Materials and Methods

### Cell lines and reagents

*Wild type* and *Fancd2*^−/−^ MEFs were grown in DMEM medium (Corning) supplemented with 10% FBS and 1% Penicillin/streptamycin (Gibco) in a 5% CO2 incubator at 37°C. RPE-1 *TP53*^−/−^ cells were grown in DMEM medium supplemented with 10% FBS and 1% Penicillin/streptamycin in a 5% CO2 incubator at 37°C. 5hmC (PY 7588; Berry&Associates) was dissolved in PBS (10 mM stock concentration). BrdU (B5002; Sigma) was dissolved in PBS and added to cell cultures at 10μM final concentration. 5-Ethynyl-2′-deoxyuridine (5-EdU) (BCN-001-100; Baseclick) and Biotin-PEG3-azide (BCFA-021-5, Baseclick) were dissolved in DMSO (10mM stock concentration). CldU (Sigma, C6891) and IdU (Sigma, I7125) were dissolved in PBS (2’5 mM stock solution), and added to cell cultures to a final concentration of 20μM and 200μM respectively. The antibodies used were against PAR (Millipore, MABE1016), ERCC1 (sc-8408), PCNA (sc-56), Lamin A/C (sc-376248), BrdU (BU1/75, Abcam), IdU (Sigma, SAB3701448), Biotin (B7653, Sigma), HMCES (NPB2-14410, Novus Biologicals), SMUG1 (E7P4I, Cell Signaling), APEX1 (WH311929, ABclonal), XRCC1 (WM038146, ABclonal), DCK (WM926071, ABclonal), DCTD (16784-1-AP, Proteintech), FLAG (F1804,Sigma), FLAG (D6W5B, Cell Signaling), PARP1 (46D11, Cell Signaling). siRNA knockdown using the D’harmacon siGENOME^TM^ siRNA pool were transfected for 24 h using RNAiMAX (Invitrogen) according to manufacturer’s instructions.

### Protein extractions and immunoblots

Cell cultures were collected by centrifugation, washed in 1 mL of ice-cold PBS and resuspended and incubated in equal volume of Benzonase buffer (2mM MgCl2, 20 mM Tris pH 8, 10% glycerol, 1% Triton X-100 and 12.5 units/mL Benzonase) on ice for 10 minutes. 2% SDS was added (1:1 vol/vol) to samples and heated at 80°C for 2 minutes. Samples were resolved on a PAGE gel, blotted onto a nitrocellulose membrane and incubated overnight with several antibodies.

### Cytotoxicity assays

2.5x10^3^ cells were seeded in 96-well plates and incubated at 37°C overnight. The day after, cells were exposed to increased doses of 5hmdC and incubated for 72 h. The assay was developed by adding the Cell Proliferation Kit I (Roche) according to the manufacturer’s instructions. Colorimetric analysis was performed using a Varioskan Flash plate reader (Thermo) at 550 nm.

### Immunofluorescence microscopy

8 x 10^4^ cells were seeded in coverslips (VWR) in 24-well plates and incubated in the presence of 5hmdC or the drug of interest for the indicated times. Cells were permeabilized using PBS + 0.25% Triton X-100 on ice for 2 min and fixed in PBS + 4% formaldehyde on ice for 15 min. Then, cells were blocked in PBS + 0.3% Tween 20 + 3% BSA for 30 min and incubated with primary antibodies overnight at 4°C. Next day, cells were washed in PBS+0.3% Tween20 three times and incubated in PBS +alexa fluor-labelled secondary antibody of chosen wavelength + 0.1mg/mL 4’6-diamidine- 2-phenylindole dihydrochloride (DAPI) for 1h at RT in dark. Finally, cells were washed in PBS + 0.3% Tween 20 three times and coverslips were mounted in Prolong® Gold Antifade Reagent (Invitrogen). For EdU immunofluorescence, cells were pulsed with EdU (10 µM) for 1h and click-it reaction (100 mM Tris-HCl pH 7.5, 1 µM Alexa488-azide, 1 mM CuSO4, 100 mM ascorbic acid) was carried out for 30 min at RT in dark conditions. At least 400 cells per condition were scored.

Images were acquired using ×40 magnification lens in Leica Thunder imager DMi8 microscope, quantified and plotted using Fiji and GraphPad Prism 9.0 software.

### SIRF assay

SIRF assay was performed as previously described by Roy and Schlacher (*79*), with few modifications. Briefly, 10^5^ cells were seeded in coverslips and the day after, EdU was added to the medium for 8 min. Cells were washed twice in PBS and medium containing 5hmdC (10 μM) was added for 3h. Cells were permeabilized in PBS + 0.5% Triton X-100 at 4°C for 2 min and fixed in PBS + 2% formaldehyde at 4°C for 15 min. PLA foci, pictures of cells were scored using 63x magnification lens in a Leica Thunder imager DMi8 microscope using LAS X software, and foci number was analyzed using a Fiji software.

### Alkaline comet assay

2·10^5^ cells were seeded in a 6-well dish and treated as indicated. After treatment, cells were collected, washed once, mixed with low melting point agarose at 1/5 ratio (v/v) and immediately layered onto pre-chilled frosted glass slides pre-coated with 0.6% agarose. Slides were allowed to settle in the dark at 4 °C for at least 30min, immersed in pre-chilled alkaline lysis buffer (2.5 M NaCl, 10 mM Tris–HCl, 100 mM EDTA, 10 mM Tris-Cl, 1% v/v DMSO, 1% v/v Triton X-100, pH 10) at 4 °C for 1 h, and incubated in pre-chilled alkaline electrophoresis buffer (1 mM EDTA, 50 mM NaOH, 1% v/v DMSO) for 30 min prior to electrophoresis at 0.6 V/cm for 25 min at 4 °C. Following electrophoresis, slides were neutralized (0.4 M Tris-HCl pH 7 for 1 h), stained with 1x SYBR green (Sigma S9430) for 10 minutes in PBS and visualized using a 10x magnification lens in a Leica Thunder imager DMi8 microscope using LAS X software. Values represent tail moment of comet and analysed by OpenComet software.

### Chromosome aberrations assays

Chromosome aberrations test was performed as previously described by Peña-Gomez et al (*15*).

### Sister chromatid exchange assays

Cells were grown in the presence of 10 μM BrdU during two cell-cycles and 5hmdC was added during the last 12 h. Chromosome were stained in 0.1M phosphate buffer solution (pH 6.8) containing 10 μg/mL Hoeschst for 25 min. Slides were washed in McIlvaine’s solution (164 mM Na2HPO4, 15 mM citric acid) and exposed to UV light (365 nm) for 1h. After that, slides were incubated in 2XSSC solution (300 mM NaCl, 30 mM Na citrate) at 62°C for 1h, and counterstained in phosphate buffer (pH 6.8) + 0.05% Tween20 + Leishman (1:4, vol:vol) for 3 min. To score chromatid exchanges, pictures of metaphases were scored using a ×100 magnification lens Leica DM6000 optical microscope and analyzed by LAS AF Leica software. Pictures were blinded- scored.

### DNA fibres

For replication fork blockage purpose, 7.5 × 10^4^ cells were seeded and pulsed with CldU (20 μM) followed by IdU (200 μM) in the presence of 5hmdC for further 30 min. A 2 μL cell drop was placed onto slides, incubated for 6 min, mixed with 7 μL spreading buffer for further 30 min at RT.

HCl (2.5 M) for 1 h at RT. Upon washing and blocking, slides were incubated with primary and secondary antibodies. Finally, slides were mounted in Fluoromount-G (eBioscience), scored with a ×40 magnification lens Leica DM6000 optical microscope using LAS AF Leica software and analyzed using a Fiji software.

### S1 nuclease fiber assay

S1 nuclease fiber assay was performed as previously described by Quinet et al. (*47*) with few modifications. Cells were incubated with CSK100 buffer (100 mM NaCl, 10 mM MOPS, pH 7, 3 mM MgCl2 pH 7.2, 300 mM sucrose, 0.5% Triton X-100) for 5 min, washed with PBS and treated with 20 U/ml S1 nuclease (ThermoFisher #EN0321) for 30 min at 37°C in S1 nuclease Buffer (30 mM sodium acetate, 10 mM zinc acetate, 5% glycerol, 50 mM NaCl, pH 4.6). Then, cells were washed with PBS and harvest with scraper.

### CRISPR/Cas9-mediated gene disruptions in RPE-1 cell lines

sgRNAs targeting exon 2 of human FANCD2 gene was previously reported (*15*). sgRNAs targeting exon 3 of HMCES gene were HMCES-3.3: 5’-AUCAUUGCUCCCAUGCGCUG-3’; HMCES-3.8: 5’- UACGGUAUCACUACGACAGU -3’. DNA oligos containing targeted sequences were cloned into pSpCas9(BB)-2A-GFP (pX458, a gift from Dr. Feng Zhang, Addgene plasmid # 48138). After transfection, single GFP^+^ sorted cells were seeded and individual clones were analyzed for HMCES deletions by PCR, further confirmed by Sanger sequencing and western blotting.

### Statistical analyses

Central line of PAR intensity signal or HMCES-FLAG nuclear intensity plots represents median value, whereas in EdU-PAR foci per cell or alkaline comet assay plots represents mean value ± s.d. Central line and boxes in DNA fiber assays represent median and 10-90 percentile. Statistical analyses were completed using Prism 8 (GraphPad). An ANOVA test was used when comparing more than two groups followed by a Tukey or Fisher’s LSD post-test. No statistical methods or criteria were used to estimate sample size or to include/exclude samples. Unless otherwise stated, all experiments were performed three times.

## Acknowledgments

RPE1 p53-/- cell line was kindly provided by Dr. Pablo Huertas (University of Seville, Spain), and Crossańs and Ponteĺs lab for critical reading of the manuscript.

## Author contributions

Conceptualization: IVR

Methodology: MJPG, MdRO

Investigation: MJPG, YRM, MdRO

Visualization: MJPG, YRM, MdRO

Supervision: IVR

Writing—original draft: IVR

Writing—review & editing: JYM, IVR

## Data and materials availability

All data analysis is available in the main text or the supplementary materials. Raw data and materials used are available to any researcher on reasonable request.

## Supplementary Materials

**Supplementary figure 1:**
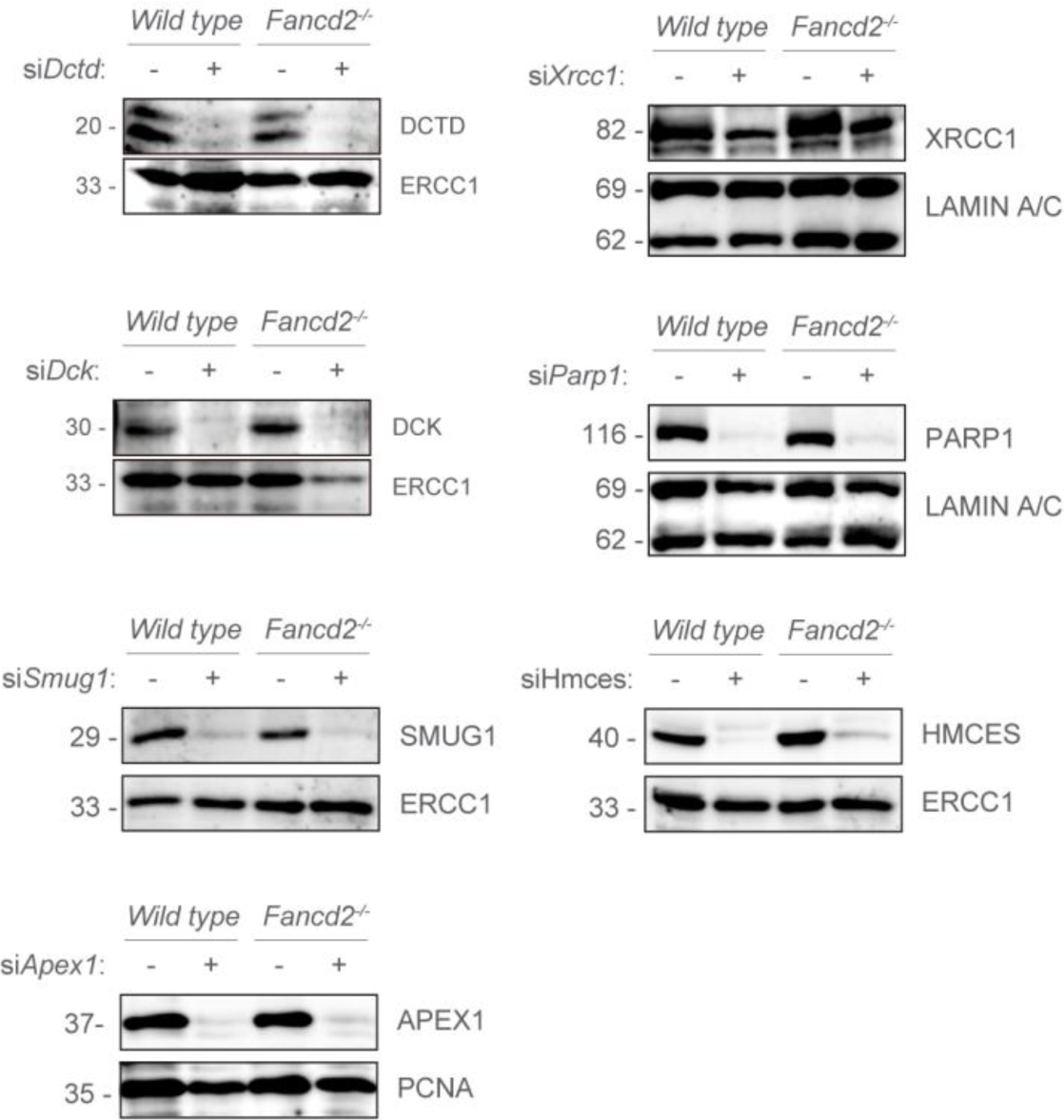
SiRNA protein depletion of DCTD, DCK, SMUG1, APEX1, XRCC1, PARP1 or HMCES examined by western blots of *wild type* and *Fancd2^-/-^* cell extracts. ERCC1, LAMIN A/C or PCNA were used as loading controls.

**Supplementary figure 2.**
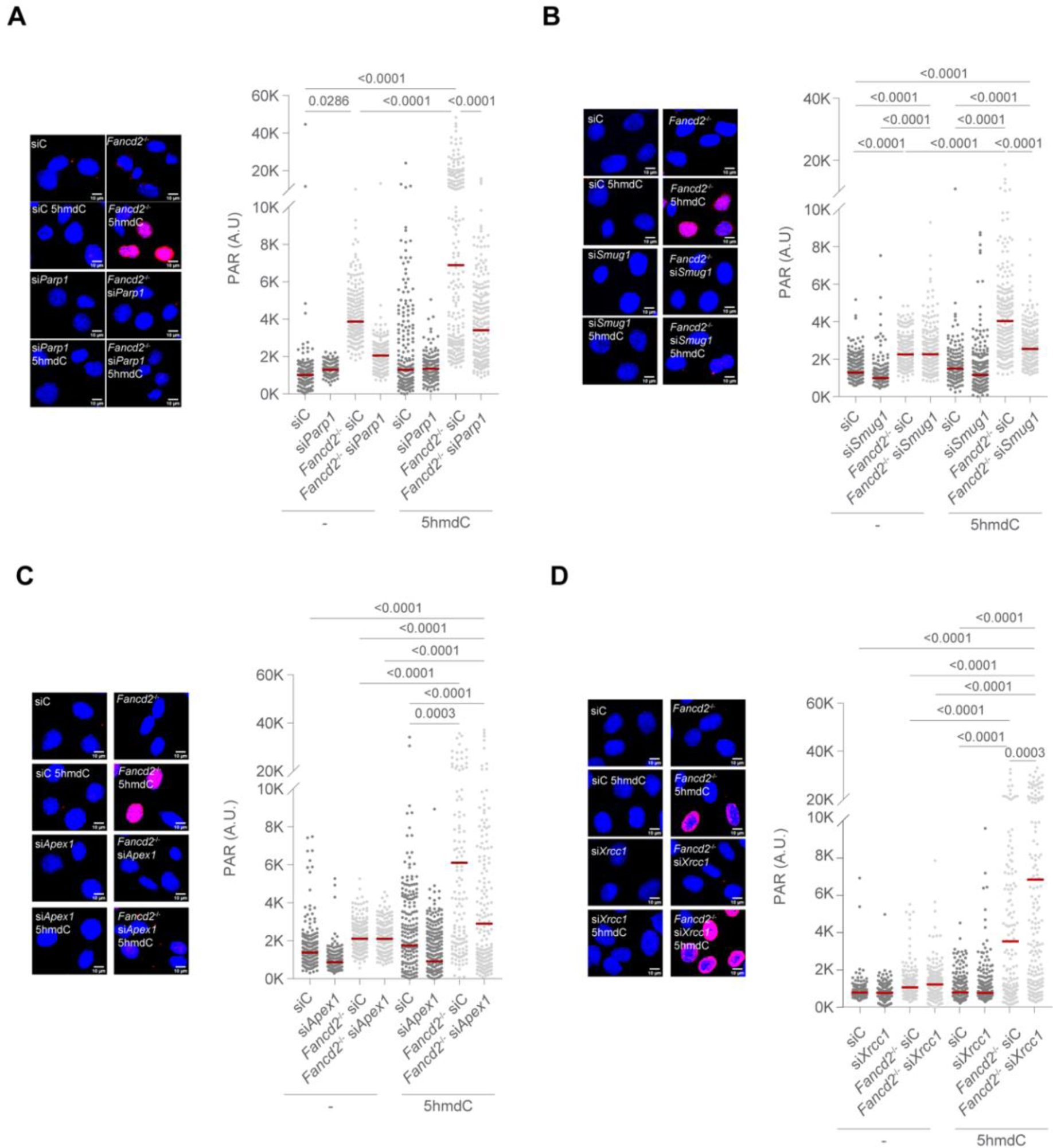
Nuclear PAR levels upon PARP1, SMUG1, APEX1 and XRCC1, PARP1 depletion of *wild type* and *Fancd2*^-/-^ cells. A. *Left*, representative total PARylation (red) immunofluorescence images of PARP1-depleted *wild type* and *Fancd2^-/-^* cells exposed to 5hmdC (10µM) for 3h. DAPI (blue) stains nuclear DNA. *Right*, plot depicting PAR mean intensity signal per nucleus. B. *Left*, representative total PARylation (red) immunofluorescence images of SMUG1-depleted *wild type* or *Fancd2^-/-^* cells and exposed to 5hmdC (10µM) for 3h. DAPI (blue) stains nuclear DNA. *Right*, plot depicting PAR mean intensity signal per nucleus. C. *Left*, representative total PARylation (red) immunofluorescence images of APEX1-depleted *wild type* or *Fancd2*^-/-^ cells exposed to 5hmdC (10µM) for 3h. DAPI (blue) stains nuclear DNA. *Right*, plot depicting PAR mean intensity signal per nucleus. D. *Left*, representative total PARylation (red) immunofluorescence images of XRCC1-depleted *wild type* or *Fancd2*^-/-^ cells exposed to 5hmdC (10µM) for 3h. DAPI (blue) stains nuclear DNA. *Right*, plot depicting PAR mean intensity signal per nucleus

**Supplementary figure 3:**
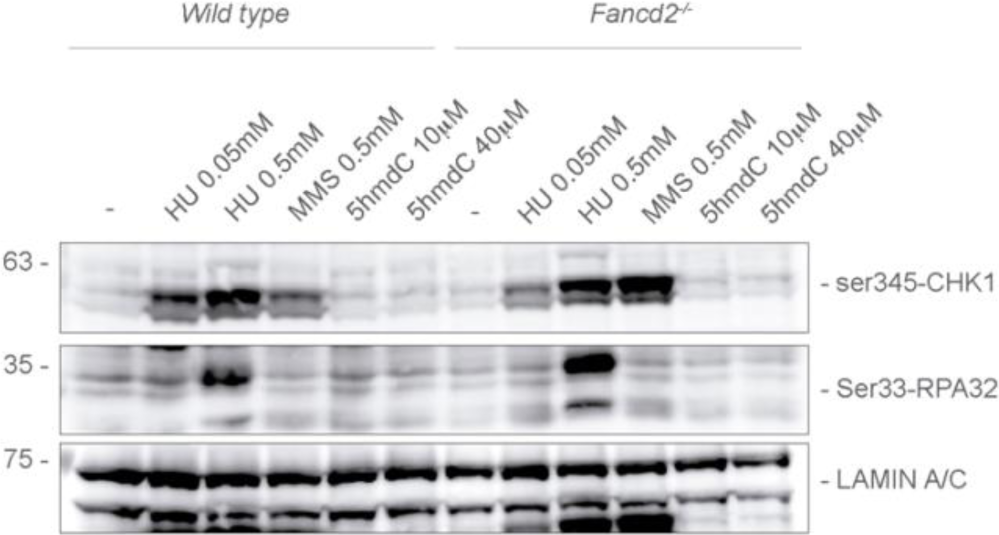
Western blots of *wild type* and *Fancd2^-/-^* cell extracts upon HU, MMS or 5hmdC treatments (3h each) to detect ser345-CHK1 or ser33-RPA2. LAMIN A/C was used as loading control.

**Supplementary figure 4.**
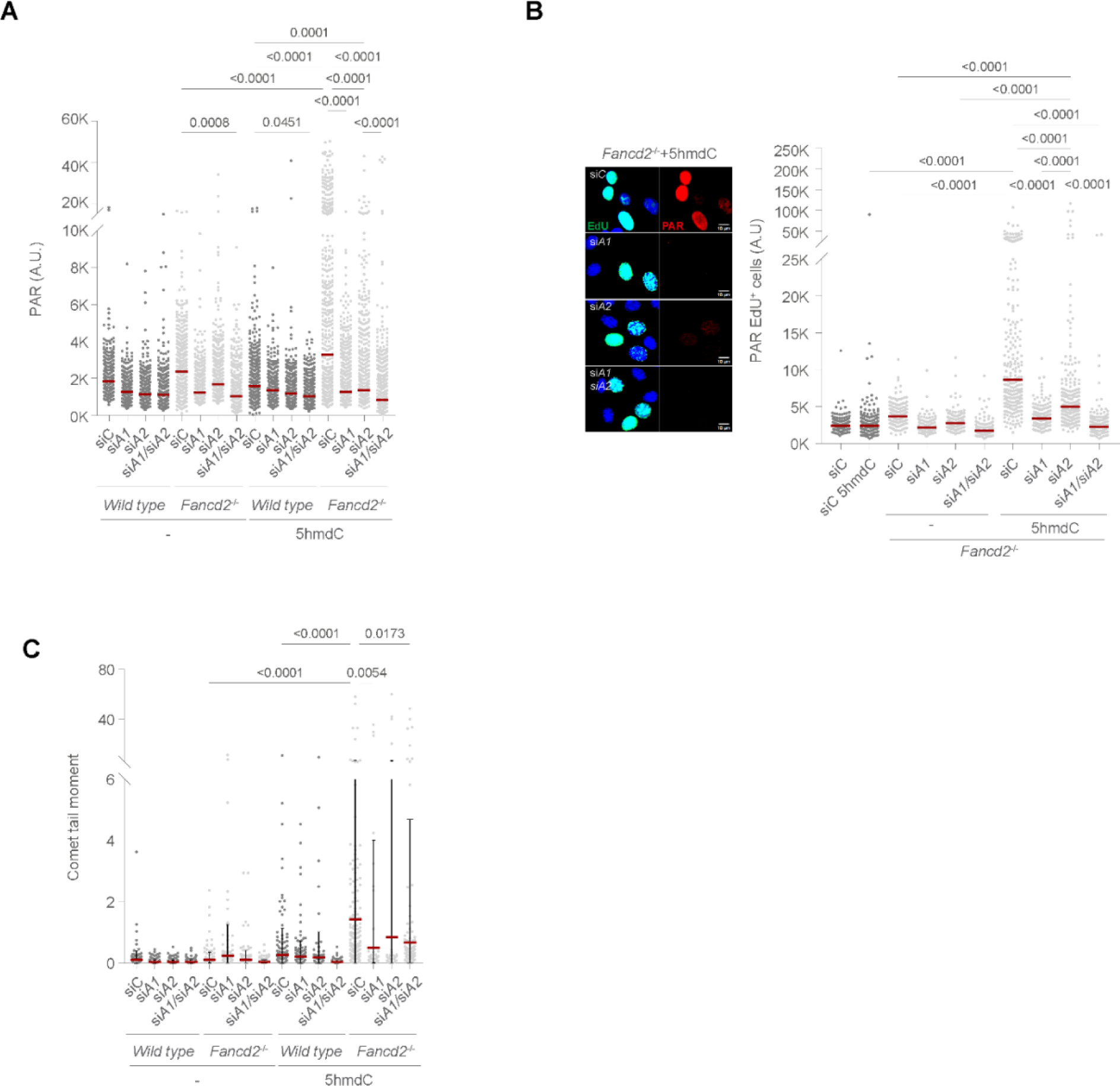
APEX1 and APEX2 contribute to 5hmdC-induced nuclear PARylation and SSBs generation in HRD cells A. Plot depicting PAR mean intensity signal per nucleus of APEX1- (siA1), APEX2- (siA2) or APEX1 APEX2 (siA1siA2)-depleted *wild type* or *Fancd2*^-/-^ cells exposed to 5hmdC (10µM). B. *Left*, representative PAR (red) immunofluorescence images of APEX1- (siA1), APEX2- (siA2) or APEX1 APEX2 (siA1siA2)-depleted *Fancd2*^-/-^ EdU^+^ cells exposed to 5hmdC (10µM) for 3h. DAPI (blue) stains nuclear DNA. EdU (green) stains S-phase cells. *Right*, plot depicting PAR mean intensity signal per nucleus. C. Plot depicting comet tail moment per cell from APEX1- (siA1), APEX2- (siA2) or APEX1 APEX2 (siA1siA2)-depleted *wild type* or *Fancd2*^-/-^ cells exposed to 5hmdC (10µM) for 3h (n=2).

**Supplementary figure 5.**
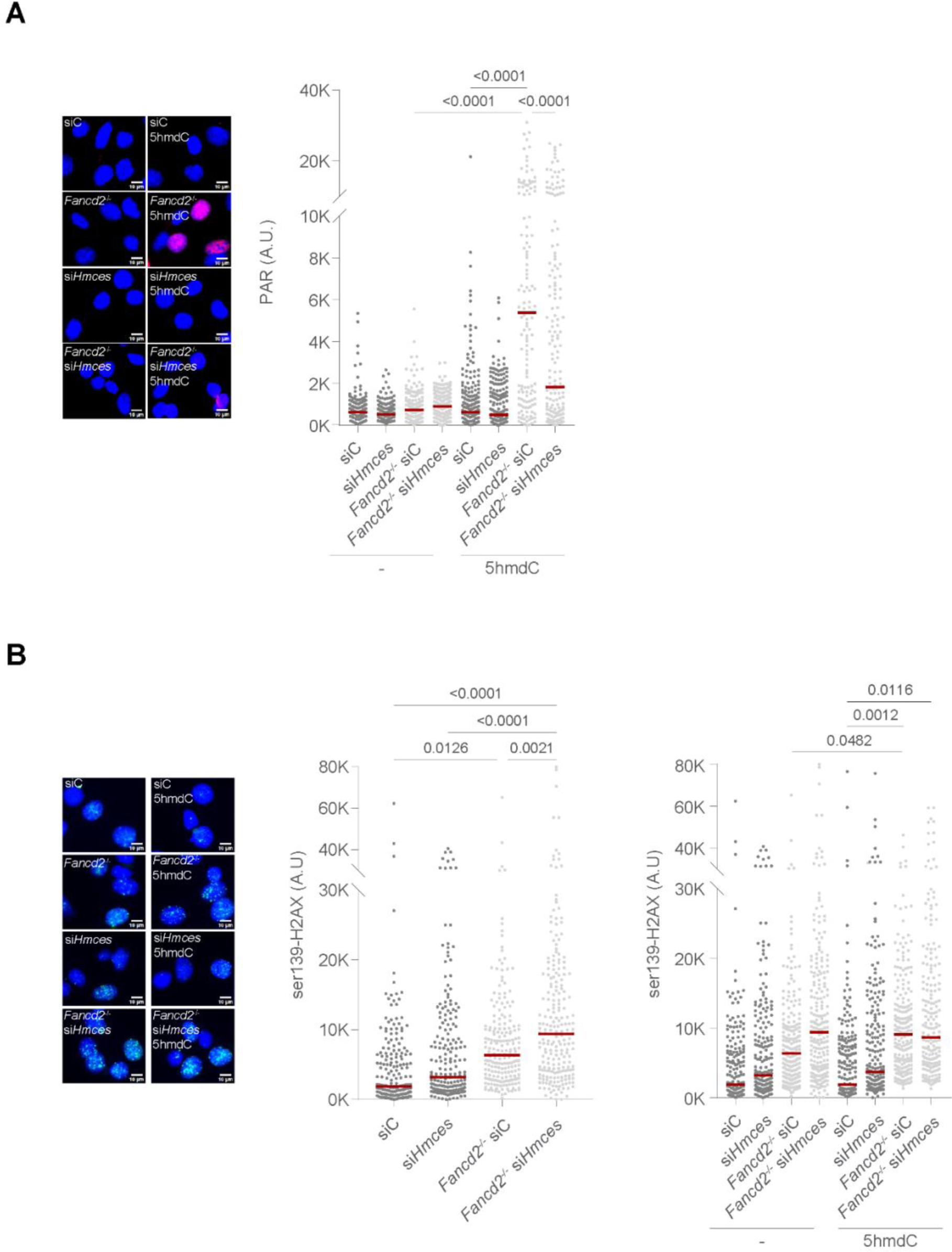
HMCES knockdown suppresses nuclear PARylation but increases DSB signalling in HRD cells A. *Left*, representative total PARylation (red) immunofluorescence images of *wild type*, *Fancd2^-/-^*, si*Hmces* and *Fancd2^-/-^* si*Hmces* cells exposed to 5hmdC (10µM) for 3h. DAPI (blue) stains nuclear DNA. *Right*, plot depicting PAR mean intensity signal per nucleus. B. *Left*, representative ser139-*Middle*, plot depicting ser139-H2AX mean intensity signal of *wild type* or *Fancd2*^-/-^ cells. *Right*, plot depicting ser139-H2AX mean intensity signal of *wild type* or *Fancd2*^-/-^ cells exposed to 5hmdC (10µM) for 3h.

**Supplementary figure 6.**
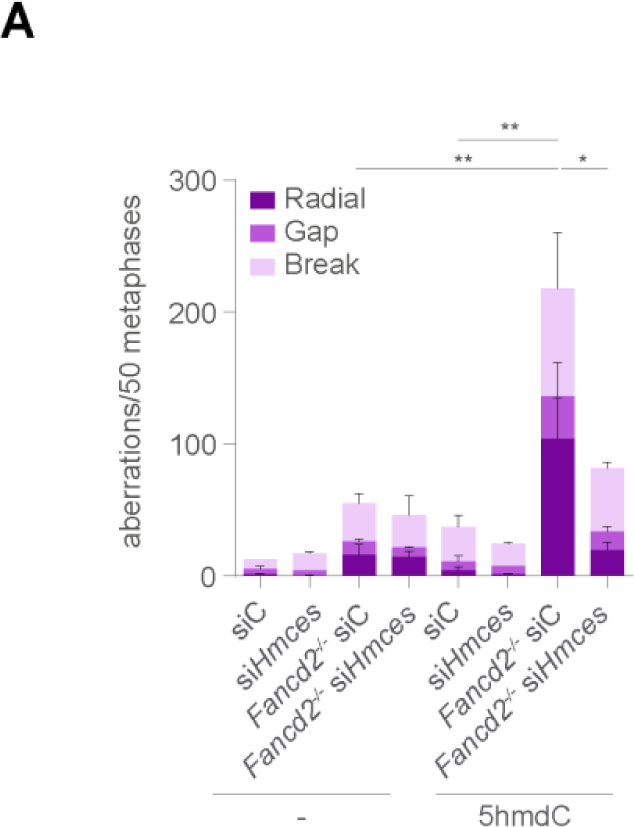
Chromosome aberrations Bar plot of different types of chromosome aberrations (Radial, gaps and breaks) from HMCES- knockdown *wild type* or *Fancd2*^-/-^ cells following 5hmdC treatment (10µM) for 40h. (n=50 of each of 2 biological replicates; bar represents mean ± s.d.).

**Supplementary figure 7.**
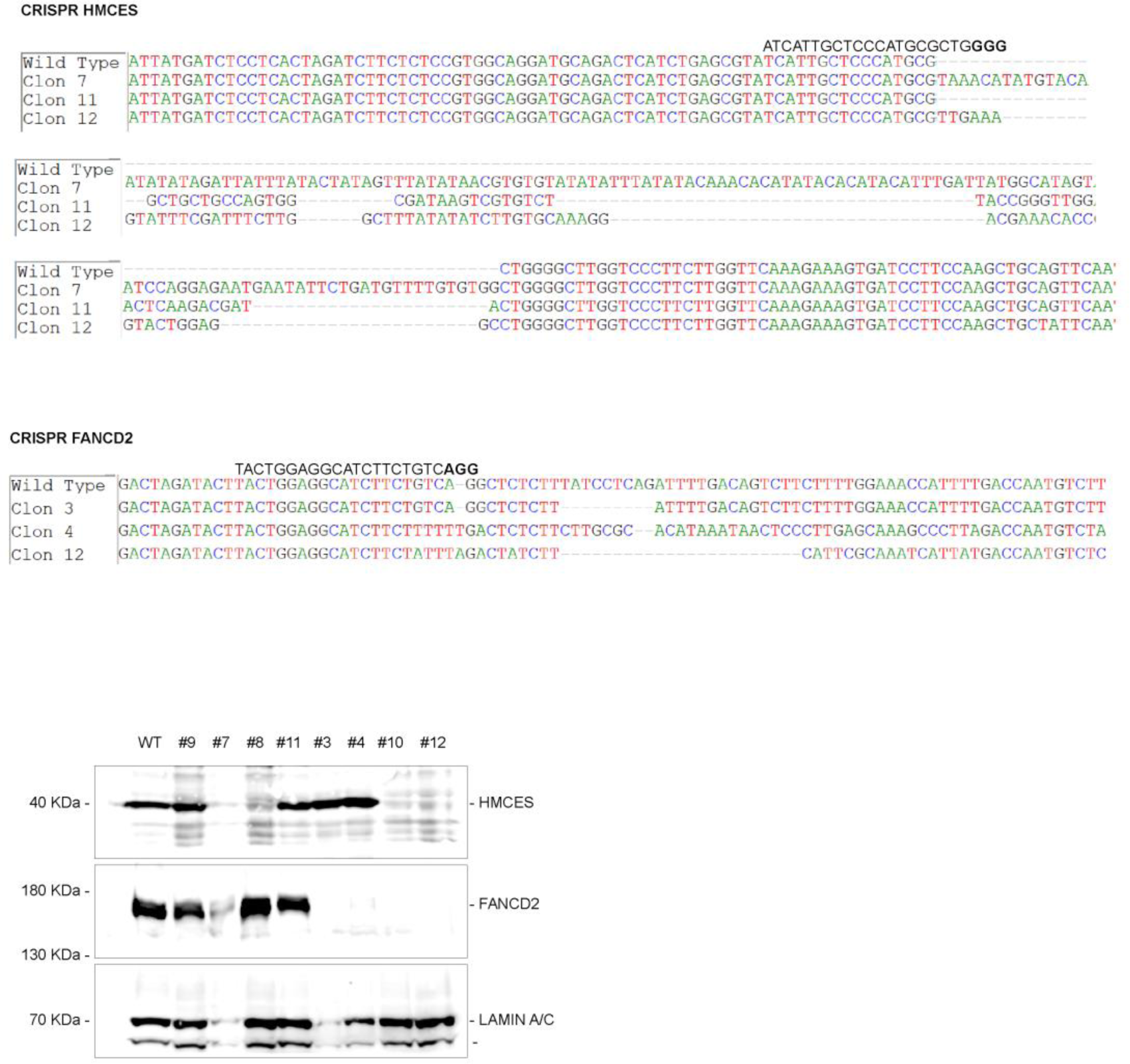
Sanger sequencing of HMCES-, FANCD2- or HMCES FANCD2 CRISPR knockout RPE-1 *TP53*^-/-^ clones. Western blot of clones obtained. Clones 9, 7, 3 and 12 were selected as *Wild type*, *HMCES*^-/-^, *FANCD2*^-/-^ and *HMCES*^-/-^*FANCD2*^-/-^ DKO respectively for further experiments.

**Supplementary figure 8.**
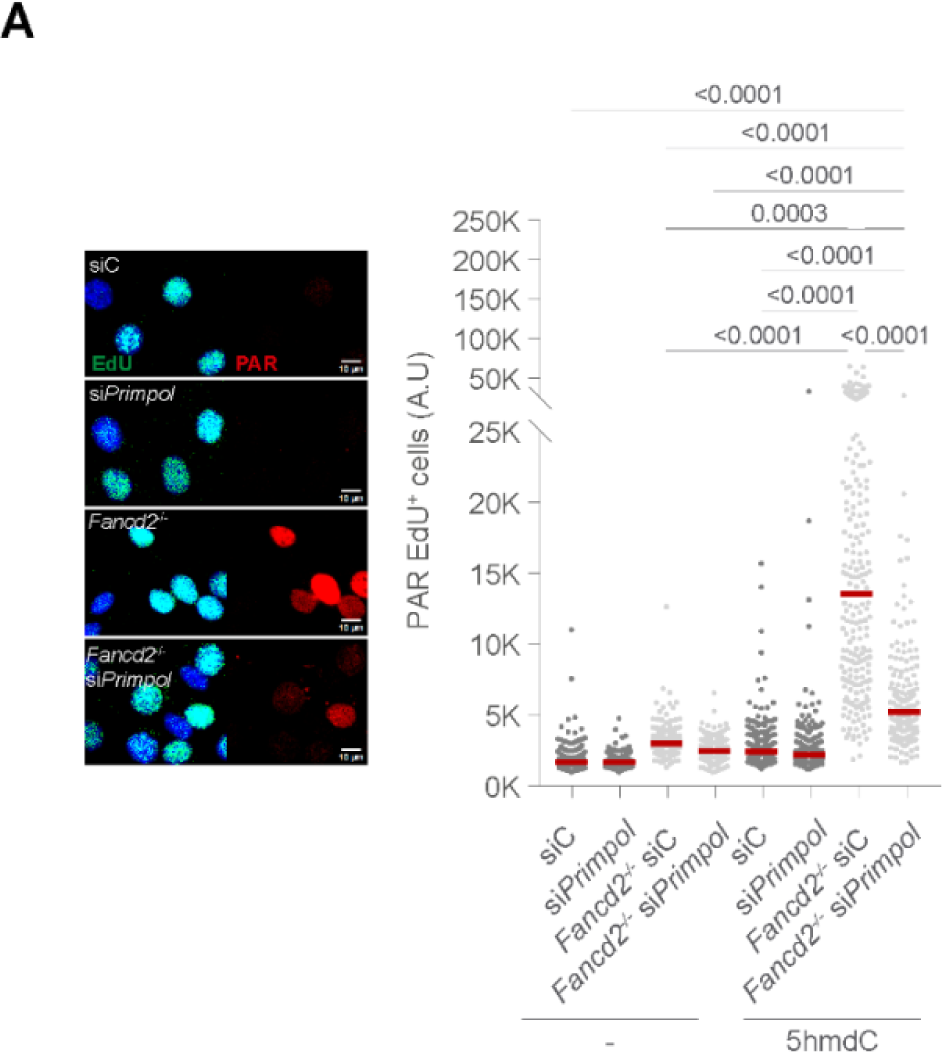
A. Plot depicting PAR mean intensity signal per PRIMPOL-depleted *wild type* or *Fancd2*^-/-^ EdU^+^ cells exposed to 5hmdC (10µM).

## References

1. M. Jung, A. Smogorzewska, Endogenous formaldehyde destroys blood stem cells. Blood 137, 1988–1990 (2021).

2. J. I. Garaycoechea et al., Genotoxic consequences of endogenous aldehydes on mouse haematopoietic stem cell function. Nature 489, 571–575 (2012).

3. A. D. Auerbach, Fanconi anemia and its diagnosis. Mutat Res 668, 4–10 (2009).

4. A. L. H. Webster et al., Genomic signature of Fanconi anaemia DNA repair pathway deficiency in cancer. Nature 612, 495–502 (2022).

5. P. R. Andreassen, A. D. D’Andrea, T. Taniguchi, ATR couples FANCD2 monoubiquitination to the DNA-damage response. Genes Dev 18, 1958–1963 (2004).

6. S. Shakeel et al., Structure of the Fanconi anaemia monoubiquitin ligase complex. Nature 575, 234–237 (2019).

7. M. C. Kottemann, A. Smogorzewska, Fanconi anaemia and the repair of Watson and Crick DNA crosslinks. Nature 493, 356–363 (2013).

8. G. L. Moldovan, A. D. D’Andrea, To the rescue: the Fanconi anemia genome stability pathway salvages replication forks. Cancer Cell 22, 5–6 (2012).

9. I. V. Rosado, F. Langevin, G. P. Crossan, M. Takata, K. J. Patel, Formaldehyde catabolism is essential in cells deficient for the Fanconi anemia DNA-repair pathway. Nat Struct Mol Biol 18, 1432–1434 (2011).

10. L. B. Pontel et al., Endogenous Formaldehyde Is a Hematopoietic Stem Cell Genotoxin and Metabolic Carcinogen. Mol Cell 60, 177–188 (2015).

11. M. S. Sasaki, A. Tonomura, A high susceptibility of Fanconi’s anemia to chromosome breakage by DNA cross-linking agents. Cancer Res 33, 1829–1836 (1973).

12. A. D. Auerbach, Diagnosis of fanconi anemia by diepoxybutane analysis. Curr Protoc Hum Genet **Chapter** 8, Unit 8 7 (2003).

13. N. G. Howlett et al., Biallelic inactivation of BRCA2 in Fanconi anemia. Science 297, 606–609 (2002).

14. K. Schlacher, H. Wu, M. Jasin, A distinct replication fork protection pathway connects Fanconi anemia tumor suppressors to RAD51-BRCA1/2. Cancer Cell 22, 106–116 (2012).

15. M. J. Peña-Gomez et al., FANCD2 maintains replication fork stability during misincorporation of the DNA demethylation products 5-hydroxymethyl-2’-deoxycytidine and 5-hydroxymethyl-2’-deoxyuridine. Cell Death Dis 13, 503 (2022).

16. Z. Kais et al., FANCD2 Maintains Fork Stability in BRCA1/2-Deficient Tumors and Promotes Alternative End-Joining DNA Repair. Cell Rep 15, 2488–2499 (2016).

17. H. E. Bryant et al., Specific killing of BRCA2-deficient tumours with inhibitors of poly(ADP-ribose) polymerase. Nature 434, 913–917 (2005).

18. R. Ceccaldi et al., Homologous-recombination-deficient tumours are dependent on Poltheta-mediated repair. Nature 518, 258–262 (2015).

19. K. Cong, S. B. Cantor, Exploiting replication gaps for cancer therapy. Mol Cell 82, 2363–2369 (2022).

20. E. Cybulla, A. Vindigni, Leveraging the replication stress response to optimize cancer therapy. Nat Rev Cancer 23, 6–24 (2023).

21. A. Ray Chaudhuri et al., Replication fork stability confers chemoresistance in BRCA-deficient cells. Nature 535, 382–387 (2016).

22. T. Lindahl, DNA repair enzymes. Annu Rev Biochem 51, 61–87 (1982).

23. D. E. Barnes, T. Lindahl, B. Sedgwick, DNA repair. Curr Opin Cell Biol 5, 424–433 (1993).

24. T. Lindahl, Recognition and processing of damaged DNA. J Cell Sci Suppl 19, 73–77 (1995).

25. A. Tubbs, A. Nussenzweig, Endogenous DNA Damage as a Source of Genomic Instability in Cancer. Cell 168, 644–656 (2017).

26. J. C. Fromme, G. L. Verdine, Base excision repair. Adv Protein Chem 69, 1–41 (2004).

27. K. W. Caldecott, Mammalian DNA base excision repair: Dancing in the moonlight. DNA Repair (Amst) 93, 102921 (2020).

28. M. Garcia-Diaz et al., A structural solution for the DNA polymerase lambda-dependent repair of DNA gaps with minimal homology. Mol Cell 13, 561–572 (2004).

29. P. J. McKinnon, K. W. Caldecott, DNA strand break repair and human genetic disease. Annu Rev Genomics Hum Genet 8, 37–55 (2007).

30. K. W. Caldecott, Single-strand break repair and genetic disease. Nat Rev Genet 9, 619–631 (2008).

31. J. L. Quinones, B. Demple, When DNA repair goes wrong: BER-generated DNA-protein crosslinks to oxidative lesions. DNA Repair (Amst) 44, 103–109 (2016).

32. S. H. Wilson, T. A. Kunkel, Passing the baton in base excision repair. Nat Struct Biol 7, 176–178 (2000).

33. A. Quinet, S. Tirman, E. Cybulla, A. Meroni, A. Vindigni, To skip or not to skip: choosing repriming to tolerate DNA damage. Mol Cell 81, 649–658 (2021).

34. K. J. Neelsen, M. Lopes, Replication fork reversal in eukaryotes: from dead end to dynamic response. Nat Rev Mol Cell Biol 16, 207–220 (2015).

35. S. Mouron et al., Repriming of DNA synthesis at stalled replication forks by human PrimPol. Nat Struct Mol Biol 20, 1383–1389 (2013).

36. A. Quinet et al., Translesion synthesis mechanisms depend on the nature of DNA damage in UV-irradiated human cells. Nucleic Acids Res 44, 5717–5731 (2016).

37. A. L. Piberger et al., PrimPol-dependent single-stranded gap formation mediates homologous recombination at bulky DNA adducts. Nat Commun 11, 5863 (2020).

38. A. Taglialatela et al., REV1-Polzeta maintains the viability of homologous recombination- deficient cancer cells through mutagenic repair of PRIMPOL-dependent ssDNA gaps. Mol Cell 81, 4008–4025 e4007 (2021).

39. A. Mann et al., POLtheta prevents MRE11-NBS1-CtIP-dependent fork breakage in the absence of BRCA2/RAD51 by filling lagging-strand gaps. Mol Cell 82, 4218–4231 e4218 (2022).

40. O. Belan et al., POLQ seals post-replicative ssDNA gaps to maintain genome stability in BRCA-deficient cancer cells. Mol Cell 82, 4664–4680 e4669 (2022).

41. H. Farmer et al., Targeting the DNA repair defect in BRCA mutant cells as a therapeutic strategy. Nature 434, 917–921 (2005).

42. K. Cong et al., Replication gaps are a key determinant of PARP inhibitor synthetic lethality with BRCA deficiency. Mol Cell 81, 3128–3144 e3127 (2021).

43. P. X. Lim, M. Zaman, W. Feng, M. Jasin, BRCA2 promotes genomic integrity and therapy resistance primarily through its role in homology-directed repair. Mol Cell 84, 447–462 e410 (2024).

44. M. J. Peña-Gomez, M. Suarez-Pizarro, I. V. Rosado, XRCC1 Prevents Replication Fork Instability during Misincorporation of the DNA Demethylation Bases 5-Hydroxymethyl- 2’-Deoxycytidine and 5-Hydroxymethyl-2’-Deoxyuridine. Int J Mol Sci 23, (2022).

45. K. Fugger et al., Targeting the nucleotide salvage factor DNPH1 sensitizes BRCA- deficient cells to PARP inhibitors. Science 372, 156–165 (2021).

46. S. Garcia-Gomez et al., PrimPol, an archaic primase/polymerase operating in human cells. Mol Cell 52, 541–553 (2013).

47. A. Quinet et al., PRIMPOL-Mediated Adaptive Response Suppresses Replication Fork Reversal in BRCA-Deficient Cells. Mol Cell 77, 461–474 e469 (2020).

48. A. Serrano-Benitez et al., Unrepaired base excision repair intermediates in template DNA strands trigger replication fork collapse and PARP inhibitor sensitivity. EMBO J 42, e113190 (2023).

49. S. Saxena et al., Unprocessed genomic uracil as a source of DNA replication stress in cancer cells. Mol Cell 84, 2036–2052 e2037 (2024).

50. K. N. Mohni et al., HMCES Maintains Genome Integrity by Shielding Abasic Sites in Single-Strand DNA. Cell 176, 144–153 e113 (2019).

51. K. P. M. Mehta, C. A. Lovejoy, R. Zhao, D. R. Heintzman, D. Cortez, HMCES Maintains Replication Fork Progression and Prevents Double-Strand Breaks in Response to APOBEC Deamination and Abasic Site Formation. Cell Rep 31, 107705 (2020).

52. L. Halabelian et al., Structural basis of HMCES interactions with abasic DNA and multivalent substrate recognition. Nat Struct Mol Biol 26, 607–612 (2019).

53. M. Srivastava et al., HMCES safeguards replication from oxidative stress and ensures error-free repair. EMBO Rep 21, e49123 (2020).

54. D. Yaneva et al., The FANCJ helicase unfolds DNA-protein crosslinks to promote their repair. Mol Cell 83, 43–56 e10 (2023).

55. M. Donsbach et al., A non-proteolytic release mechanism for HMCES-DNA-protein crosslinks. EMBO J 42, e113360 (2023).

56. J. Rua-Fernandez et al., Self-reversal facilitates the resolution of HMCES-DNA protein crosslinks in cells. bioRxiv, (2023).

57. H. Hanzlikova et al., The Importance of Poly(ADP-Ribose) Polymerase as a Sensor of Unligated Okazaki Fragments during DNA Replication. Mol Cell 71, 319–331 e313 (2018).

58. A. Vaitsiankova et al., PARP inhibition impedes the maturation of nascent DNA strands during DNA replication. Nat Struct Mol Biol 29, 329–338 (2022).

59. A. Taglialatela et al., Restoration of Replication Fork Stability in BRCA1- and BRCA2- Deficient Cells by Inactivation of SNF2-Family Fork Remodelers. Mol Cell 68, 414–430 e418 (2017).

60. T. C. Liu et al., APE1 distinguishes DNA substrates in exonucleolytic cleavage by induced space-filling. Nat Commun 12, 601 (2021).

61. P. Burkovics, I. Hajdu, V. Szukacsov, I. Unk, L. Haracska, Role of PCNA-dependent stimulation of 3’-phosphodiesterase and 3’-5’ exonuclease activities of human Ape2 in repair of oxidative DNA damage. Nucleic Acids Res 37, 4247–4255 (2009).

62. A. Alvarez-Quilon et al., Endogenous DNA 3’ Blocks Are Vulnerabilities for BRCA1 and BRCA2 Deficiency and Are Reversed by the APE2 Nuclease. Mol Cell 78, 1152–1165 e1158 (2020).

63. K. E. Mengwasser et al., Genetic Screens Reveal FEN1 and APEX2 as BRCA2 Synthetic Lethal Targets. Mol Cell 73, 885–899 e886 (2019).

64. A. A. Demin et al., XRCC1 prevents toxic PARP1 trapping during DNA base excision repair. Mol Cell 81, 3018–3030 e3015 (2021).

65. M. A. Hossain et al., APE2 Is a General Regulator of the ATR-Chk1 DNA Damage Response Pathway to Maintain Genome Integrity in Pancreatic Cancer Cells. Front Cell Dev Biol 9, 738502 (2021).

66. Y. Lin et al., APE1 recruits ATRIP to ssDNA in an RPA-dependent and -independent manner to promote the ATR DNA damage response. Elife 12, (2023).

67. J. Biayna et al., Loss of the abasic site sensor HMCES is synthetic lethal with the activity of the APOBEC3A cytosine deaminase in cancer cells. PLoS Biol 19, e3001176 (2021).

68. T. Lindahl, An N-glycosidase from Escherichia coli that releases free uracil from DNA containing deaminated cytosine residues. Proc Natl Acad Sci U S A 71, 3649–3653 (1974).

69. E. Seeberg, L. Eide, M. Bjoras, The base excision repair pathway. Trends Biochem Sci 20, 391–397 (1995).

70. Y. Lin et al., APE1 senses DNA single-strand breaks for repair and signaling. Nucleic Acids Res 48, 1925–1940 (2020).

71. A. Brambati et al., RHINO directs MMEJ to repair DNA breaks in mitosis. Science 381, 653–660 (2023).

72. Y. Sugimoto et al., Novel mechanisms for the removal of strong replication-blocking HMCES- and thiazolidine-DNA adducts in humans. Nucleic Acids Res 51, 4959–4981 (2023).

73. V. Shukla et al., HMCES Functions in the Alternative End-Joining Pathway of the DNA DSB Repair during Class Switch Recombination in B Cells. Mol Cell 77, 384–394 e384 (2020).

74. C. Gelot et al., Poltheta is phosphorylated by PLK1 to repair double-strand breaks in mitosis. Nature 621, 415–422 (2023).

75. J. Zhou et al., A first-in-class Polymerase Theta Inhibitor selectively targets Homologous-Recombination-Deficient Tumors. Nat Cancer 2, 598–610 (2021).

76. R. Zellweger et al., Rad51-mediated replication fork reversal is a global response to genotoxic treatments in human cells. J Cell Biol 208, 563–579 (2015).

77. A. Hale, A. Dhoonmoon, J. Straka, C. M. Nicolae, G. L. Moldovan, Multi-step processing of replication stress-derived nascent strand DNA gaps by MRE11 and EXO1 nucleases. Nat Commun 14, 6265 (2023).

78. N. Garcia-Rodriguez, I. Dominguez-Garcia, M. D. C. Dominguez-Perez, P. Huertas, EXO1 and DNA2-mediated ssDNA gap expansion is essential for ATR activation and to maintain viability in BRCA1-deficient cells. Nucleic Acids Res, (2024).

79. S. Roy, K. Schlacher, SIRF: A Single-cell Assay for in situ Protein Interaction with Nascent DNA Replication Forks. Bio Protoc 9, e3377 (2019).

